# Single-nucleus and spatial landscape of the sub-ventricular zone in human glioblastoma

**DOI:** 10.1101/2024.04.24.590852

**Authors:** Y. Licón-Muñoz, V. Avalos, S. Subramanian, B. Granger, F. Martinez, S. Varela, D. Moore, E. Perkins, M. Kogan, S. Berto, M.O. Chohan, C.A. Bowers, S.G.M. Piccirillo

## Abstract

The sub-ventricular zone (SVZ) is the most well-characterized neurogenic area in the mammalian brain. We previously showed that in 65% of patients with glioblastoma (GBM), the SVZ is a reservoir of cancer stem-like cells that contribute to treatment resistance and emergence of recurrence. Here, we built a single-nucleus RNA-sequencing-based microenvironment landscape of the tumor mass (T_Mass) and the SVZ (T_SVZ) of 15 GBM patients and 2 histologically normal SVZ (N_SVZ) samples as controls. We identified a mesenchymal signature in the T_SVZ of GBM patients: tumor cells from the T_SVZ relied on the *ZEB1* regulatory network, whereas tumor cells in the T_Mass relied on the *TEAD1* regulatory network. Moreover, the T_SVZ microenvironment was predominantly characterized by tumor-supportive microglia, which spatially co-exist and establish heterotypic interactions with tumor cells. Lastly, differential gene expression analyses, predictions of ligand-receptor and incoming/outgoing interactions, and functional assays revealed that the IL-1β/IL-1RAcP and Wnt-5a/Frizzled-3 pathways are therapeutic targets in the T_SVZ microenvironment. Our data provide insights into the biology of the SVZ in GBM patients and identify specific targets of this microenvironment.

## Main text

Glioblastoma (GBM) is a fatal disease of the adult Central Nervous System. Poor survival and extensive heterogeneity leading to treatment resistance and emergence of the recurrent tumor are key clinical and biological features. Clinical management of GBM is challenging due to its heterogeneous nature, invasive potential, and poor response to radio- and chemo-therapy^1,2^. As a result, GBM inevitably recurs^2^ and only 6.9% of patients survive 5 years post-diagnosis^3^. In 2015, we were the first to show that in most patients with GBM, the sub-ventricular zone (SVZ) of the lateral ventricles is a reservoir of cancer stem-like cells (CSCs) that show distinct patterns of treatment resistance compared to matched CSCs from the tumor mass, and contribute to seeding of the recurrent tumor^4,5^. Despite the extensive inter-tumor heterogeneity within GBM, in nearly 80% of patients, the SVZ classifies as the molecular subtype with the worst prognosis^4^, characterized by the presence of tumor-associated macrophages (TAMs)^6–28^, which consist of monocyte-derived macrophages (MDM) and microglia. Therefore, identifying therapeutic targets in the SVZ is key to developing more effective treatments. However, sampling and characterization of the SVZ in GBM is challenging, as this area is extremely small and must be objectively identified during tumor surgical resection.

Given the cellular and molecular intra-tumor heterogeneity characteristic of GBM, the functional role of tumor, stromal, and immune cells in the SVZ cannot be predicted based on analyses of samples from the tumor mass. To overcome this challenge, using our fluorescence-guided multiple sampling (FGMS) scheme^29^, we built a single-nucleus RNA-sequencing (sn-RNAseq)-based microenvironment landscape of the SVZ (T_SVZ) using tissues from 15 GBM patients (14 IDH IDH wild-type, 1 IDH mutant). For 14 of these patients, we circumvented the limitations of single-cell RNAseq by using sn-RNAseq, which allowed us to include frozen samples and preserve the cell composition in the tumor microenvironment. By systematically comparing the T_SVZ with tumor mass (T_Mass) samples isolated from the same patients and two histologically normal SVZ (N_SVZ) samples, and using a number of computational tools and experimental methods, we identified two pathways that represent novel targets in the T_SVZ microenvironment.

### A single-nucleus landscape of the tumor mass, tumor SVZ, and normal SVZ microenvironments in GBM patients

Using our FGMS scheme^29^, we built a single-nucleus landscape of the T_Mass, T_SVZ, and N_SVZ microenvironments in GBM. We collected 15 T_Mass samples and 15 matched T_SVZ samples from 15 untreated patients undergoing surgical resection for presumed high-grade glioma (**Suppl. Table 1** summarizes patient clinical and molecular information). For each sample, we defined the status of the GBM genetic drivers^30,31^ (**Suppl. Table 2**). As controls, 2 N_SVZ samples were collected from two individuals: one SVZ was collected postmortem and the other during tumor surgical resection. We performed sn-RNAseq using gel bead-in-emulsion technology. We obtained 6.3×10^6^ nuclei from T_Mass samples, 8.0×10^6^ nuclei from T_SVZ samples, and 1.1×10^6^ nuclei from N_SVZ samples (**Fig. 1a** and **Suppl. Table 3**). For each patient and each area (T_Mass, T_SVZ, and N_SVZ), we determined the number of detected genes and unique molecular identifiers (**Suppl. Fig. 1**). We sequenced about 3-7×10^3^ nuclei/sample. An estimated total of 59,967, 30,223, and 7,534 cells were detected after sequencing for T_Mass, T_SVZ, and N_SVZ, respectively (**Fig. 1a**). Our pipeline includes cell type annotation to define the T_Mass, T_SVZ, and N_SVZ landscapes and subsequent bioinformatic analyses and experimental work to identify transcription factor regulatory networks, define cellular dynamics, identify differentially expressed genes of the three areas, and characterize TAMs. These steps were followed by spatial transcriptomics, ligand-receptor predictions, and functional phenotyping to identify interactions specific to the T_SVZ and define their clinical significance (**Fig. 1a**). We integrated data from all patients by areas (T_Mass, T_SVZ, and N_SVZ, **Fig. 1b top left**), and by cluster (**Fig. 1b top right**), and calculated the proportions of cells in the three areas for each patient by cluster (**Fig. 1c**). Of note, cluster 15 was exclusive to the two N_SVZ samples (histologically normal samples 1 and 2, HNS1 and 2), confirming that these two samples were distinguishable from the T_Mass and T_SVZ (**Fig. 1c**).

**Figure 1.**
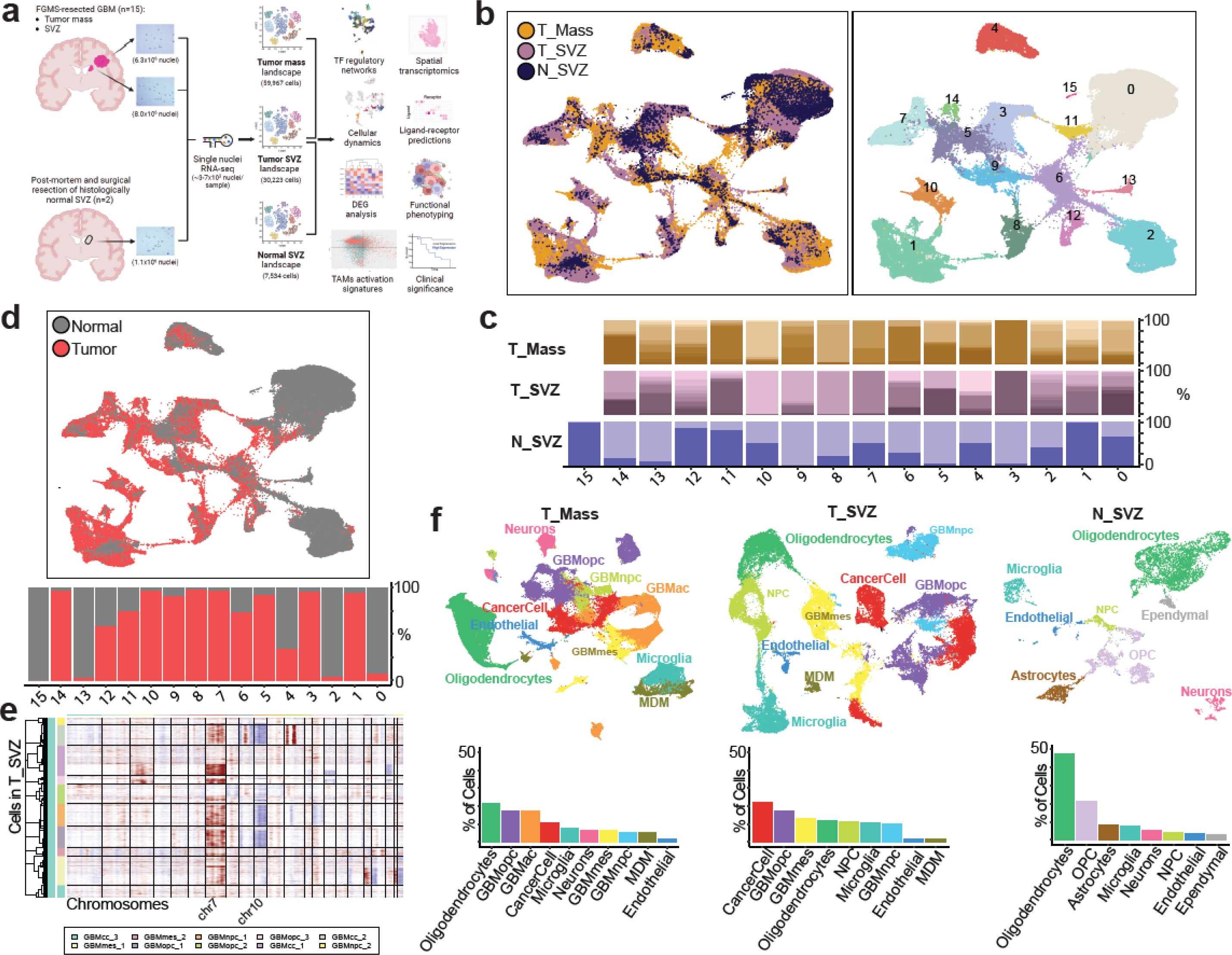
A single-nucleus landscape of the tumor mass, tumor SVZ, and normal SVZ microenvironments in GBM patients. **a,** Schematic of the tissue collection pipeline using our previously published fluorescence-guided multiple sampling (FGMS) scheme^29^. Tumor mass and SVZ samples were collected from 15 GBM patients. Histologically normal SVZ samples were collected from two individuals: one SVZ was collected as postmortem tissue and the other during tumor surgical resection. Due to poor sample quality, cells obtained from GBM4 are not included in the counting shown in this schematic. The number of cells obtained from each area, based on the number of total nuclei sequenced, is in parentheses under each illustrative Uniform Manifold Approximation and Projection (UMAP). Bioinformatic analysis and experimental work were performed to identify the transcription factor (TF) regulatory networks, define cellular dynamics, identify differentially expressed genes (DEG) and tumor-associated macrophages (TAMs) activation signatures, followed by spatial transcriptomics, ligand-receptor predictions, and functional phenotyping to identify interactions specific of the tumor SVZ with clinical significance. Created with BioRender.com. **b,** Integrated UMAPs of all cells by area (Tumor Mass=T_Mass, yellow; Tumor SVZ=T_SVZ, purple; Normal SVZ=N_SVZ, blue) (left) and cluster (right). **c,** Proportions of cells in the three areas from each patient by cluster (same color code as in b). **d,** Integrated UMAP of all cells colored by class (normal=grey and tumor=red based on copy-number variations, top) and proportions of cells from each class by cluster (bottom). **e,** Copy-number variations in integrated T_SVZ samples and clustered by cancer cell types (y-axis). Genomic region of each variation is presented by chromosomal location (x-axis). Reference cells are from integrated N_SVZ samples (**Suppl.** Fig. 2). **f,** UMAPs of each area showing cell type annotation. T_Mass (left), T_SVZ (middle), and N_SVZ (right), top. Proportion of each cell type in the three areas, bottom. Tumor cells were annotated using the cell state classification of Neftel^16^. NPC, neural progenitor cells; OPC, *oligodendrocyte precursor cells;* MDM, monocyte-derived macrophages.

Transcriptional analysis of copy-number variations (CNVs) by inferCNV^32^ provides a method to distinguish tumor and normal cells. We first used integrated data to identify tumor and normal cells by cluster (**Fig. 1d top**), and then calculated proportions of tumor and normal cells by clusters (**Fig. 1d bottom**). As expected, cluster 15 was composed entirely of normal cells. While we could identify expected CNVs such as chromosome 7 amplification and chromosome 10 deletion in the T_SVZ (**Fig. 1e**) and the T_Mass (**Suppl. Fig. 2**), the N_SVZ had a normal chromosomal landscape, confirming that it was composed of normal cells only (**Suppl. Fig. 2**).

For each cell subset identified by clustering in each area, we annotated cell types based on known marker genes and reference classifiers (**Fig. 1f top** and **Suppl. Fig. 3**). Tumor cells depicted in Fig. 1f were assigned using the cell state classification by Neftel *et al.*^16^. We also calculated the proportion of each cell type in the three areas (**Fig. 1f bottom**). Among the tumor cell states, the astrocyte-like (GBMac) state was not represented in the T_SVZ (**Fig. 1f bottom middle**), while tumor cells in matched T-Mass samples represent all four cell states: GBMac, oligodendrocyte-progenitor-like (GBMopc), mesenchymal-like (GBMmes), and neural-progenitor-like (GBMnpc) (**Fig. 1f bottom left**). While GBMopc and GBMac were present in similar proportions in the T_Mass (17.4% and 17.2%, respectively) and were the most abundant tumor cell populations in this area, the GBMmes was the second most abundant state in the T_SVZ (13.2% *versus* 6.5% in the T_Mass). The T_SVZ also showed an increase in the GBMnpc state (10.1% *versus* 5.5% in the T_Mass). Some tumor cell clusters in both areas (22.2% in the T_SVZ and 10.5% in the T_Mass) could not be captured by the existing four states^16^; we labeled those clusters as CancerCell. The same analysis performed for each patient and each area allowed us to quantify the abundance of each tumor cell state^16^ (**Suppl. Fig. 4**).

Among tumor-associated macrophages (TAMs), microglia were more abundant in the T_SVZ compared to MDM (10.5% *versus* 1.7%) (**Fig. 1f bottom middle**), whereas both cell types were similar in the T_Mass (7.6% and 5.5%, respectively) (**Fig. 1f bottom left**). The N_SVZ was composed of the expected normal brain cell types, including oligodendrocytes, astrocytes, and neurons. The correct sampling of the tissue adjacent to the ventricle in the N_SVZ was confirmed by the presence of ependymal cells (**Fig. 1f bottom right**), in agreement with a previous report^33^. Overall, these results show that the T_SVZ had a different cellular landscape than the T_Mass and the N_SVZ.

### The tumor SVZ microenvironment harbors tumor cell populations characterized by a ZEB1-centered mesenchymal signature and a distinct regulon profile of microglia

We started our analysis by defining the cellular dynamics of tumor cells in the T_Mass and the T_SVZ. Using Cell Rank^34,35^, we identified macrostates in the two areas: the T_Mass was characterized by three GBMac macrostates (GBMac p1, p2, and p3), one GBMopc, one GBMnpc, and one GBMmes. In contrast, the T_SVZ was characterized by three GBMmes macrostates (GBMmes p1, p2, and p3), two CancerCell (CancerCell p1, and p2), and one GBMopc (**Fig. 2a**). CancerCell and GBMmes cells were more undifferentiated than other tumor cells in the T_Mass and in the T_SVZ, respectively, suggesting different transcriptional dynamics and directional flows among cell populations (**Fig. 2a**). We then identified initial and terminal states (**Fig. 2b** and **Suppl. Fig. 5**), and fate probabilities (**Suppl. Fig. 6**) and asked whether the initial macrostates differed between the two areas. The initial macrostate of the T_Mass was GBMac p3, whereas for the T_SVZ it was GBMmes p1 (**Fig. 2b**). Moreover, latent time analysis revealed additional differences between the T_Mass (**Fig. 2c left**) and the T_SVZ (**Fig. 2c right**) at the level of transcription factors (TF) and co-factors. Only the T_SVZ had distinct expression patterns: some TF and co-factors, such as *ANXA11*, *HIF1A*, and *FOXO1*, showed an initial expression trend, whereas others, such as *GLI2*, and *ID4*, had a terminal expression trend (**Fig. 2c right**). In contrast, the T_Mass was characterized by TF and co-factors with terminal and more homogeneous expression trends (**Fig. 2c left**).

**Figure 2.**
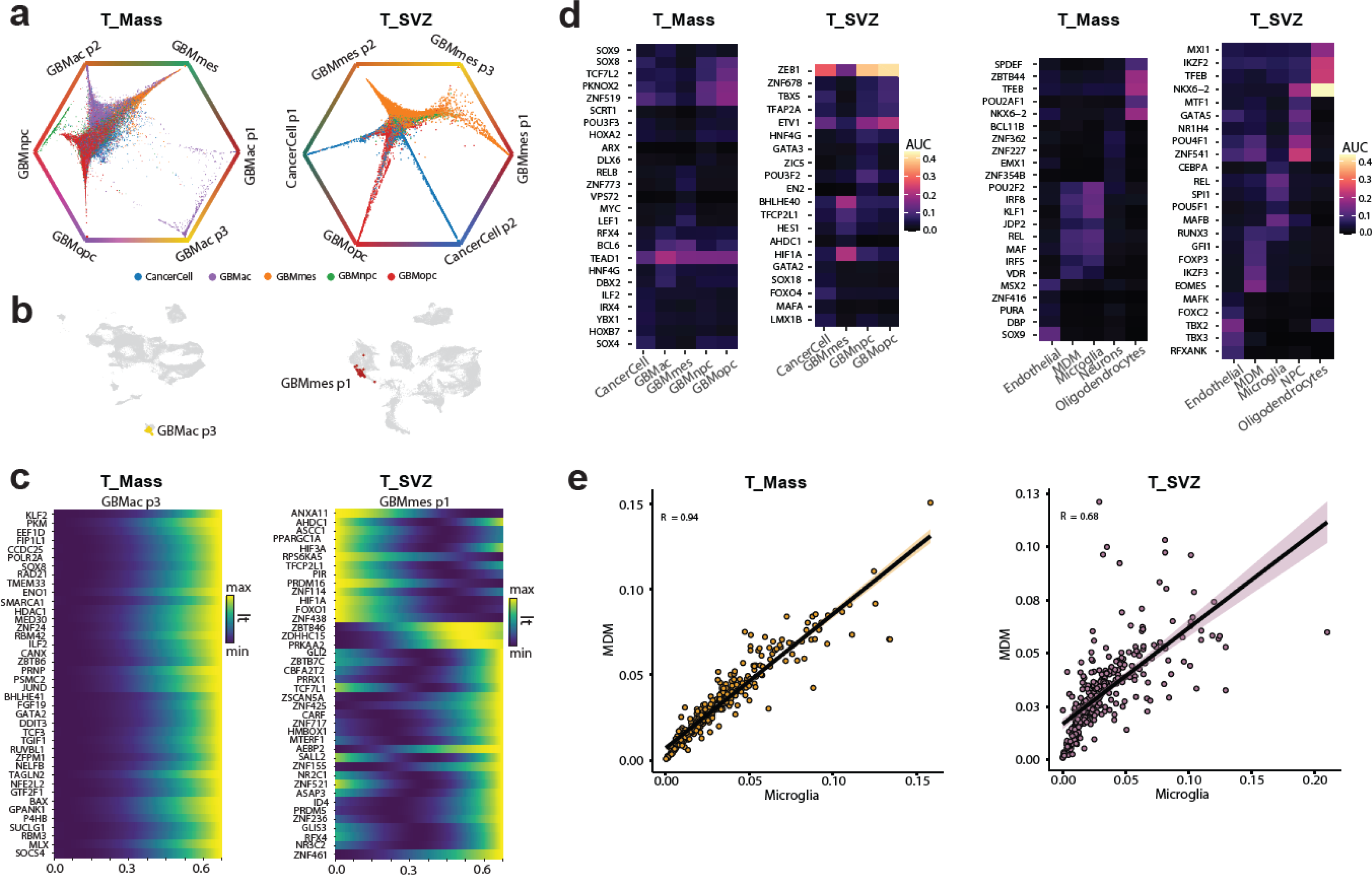
The tumor SVZ microenvironment harbors tumor cell populations characterized by a ZEB1-centered mesenchymal signature and a distinct regulon profile of microglia. **a,** Circular projections of tumor cells according to fate probabilities towards the macrostates, with cell type annotation as in Fig. 1f. Only tumor cells of the T_Mass (top left) and T_SVZ (top right) were analyzed using Cell Rank 2^34,35^. **b,** CellRank-computed initial macrostates in the T_Mass (bottom left) and T_SVZ (bottom right). Macrostates are color-coded as in Fig. 2a. **c,** Area-specific heatmaps (T_Mass, left and T_SVZ, right) showing gene expression trends of the top 40 genes with expression sorted according to latent time. Only transcription factors and co-factors are shown. **d,** Heatmaps of regulon enrichment expressed as area under the curve (AUC) in tumor cells of the T_Mass and the T_SVZ. Regulons were identified by single-cell regulatory network inference and clustering (SCENIC)^36,37^. Tumor cells of each area (left) and normal cells (right) are depicted. Only the top 5 regulons for each cell type are shown. Cell type annotation as in Fig. 1f. **e,** Pearson’s correlation analysis of regulons between monocyte-derived macrophages (MDM) and microglia in the T_Mass (left) and T_SVZ (right). *p*<2.2E-16 for both correlations.

Given the different cellular dynamics in the tumor cells of the T_SVZ compared to the T_Mass, we then performed single-cell regulatory network inference and clustering (SCENIC)^36,37^ to identify the key master regulators of the tumor cells in the T_SVZ. While the TF and regulator of cell migration *TEAD1*^38,39^ was specific to tumor cells in the T_Mass, especially in the GBMac (**Fig. 2d left**), the key mesenchymal transcription factor *ZEB1*^40–42^ was exclusively expressed in the T_SVZ, with highest enrichment in the GBMopc and GBMnpc states and lowest in the GBMmes state (**Fig. 2d left**). This may be because the GBMmes state was defined by genes related to wound healing, inflammatory response, and hypoxia, among others, but not mesenchymal processes, as recently suggested^43^. Other regulons highlighted distinct regulatory networks in the two areas: (i) *ZNF519*, *PKNOX2*, and *TCF7L2* showed enrichment in the GBMopc state of the T_Mass. Notably, *TCF7L2* is an effector of the Wnt/β-Catenin signaling pathway with prognostic significance in GBM patients^44^ (**Fig. 2d left**); (ii) *ETV1*, *BHLHE40*, and *HIF1A* were enriched in the T_SVZ, with *ETV1* having the highest enrichment in the GBMopc, whereas *BHLHE40* and *HIF1A* were highest in the GBMmes state, as expected for a state linked to hypoxia genes^16,43^ (**Fig. 2d left**).

We next calculated regulon enrichment in the matched normal cells of the T_Mass and the T_SVZ (**Fig. 2d right**). Although the T_Mass MDM and microglia shared a core set of regulons, including *POU2F2*, *KLF1*, and the regulator of macrophage differentiation *IRF8* (which confers an immunosuppressive phenotype in MDM^45^ and promotes reactivity in microglia^46^), in the T_SVZ, MDM and microglia were characterized by distinct regulons, including *REL* and *MAFB* in microglia and *EOMES* in MDM (**Fig. 2d right**), suggesting differences in the functional roles of these immune cell populations. To further explore these differences, we performed correlation analyses of regulons in MDM and microglia in the T_Mass (**Fig. 2e left**) and in the T_SVZ (**Fig. 2e right**). MDM and microglia in the T_SVZ showed a weaker correlation than the same cell populations in the T_Mass (R=0.68 *versus* R=0.94).

Overall, these results indicate that the T_SVZ harbors tumor cells characterized by a ZEB1-centered mesenchymal signature and a distinct regulon profile of microglia. Identification of a mesenchymal phenotype in the T_SVZ is consistent with our previous work in an independent cohort of SVZ samples from GBM patients^4^.

### Tumor-supportive microglia are the majority of TAMs in the T_SVZ microenvironment and spatially co-exist with tumor cells

Our results above led us to perform functional characterization of microglia. Initially, we visualized only clusters of normal (stromal and immune) cells of the T_Mass and the T_SVZ and calculated the proportion of each cell type (**Fig. 3a**). This analysis confirmed that MDM represent the minority of TAMs and the least abundant normal cell type of the T_SVZ (4.6% of total stromal/immune cells), whereas microglia were the third most abundant cell type in this area (28.4% of total stromal/immune cells; **Fig. 3a right**). In the T_Mass, microglia and MDM were present in more similar proportions (17.7% and 12.8% of total stromal/immune cells, respectively; **Fig. 3a left**). Neural progenitor cells (NPCs) represent the second most abundant cell type in the T_SVZ but were absent in the T_Mass microenvironment. Conversely, neurons represent the third most abundant cell type in the T_Mass but were absent in the T_SVZ (**Fig. 3a**).

**Figure 3.**
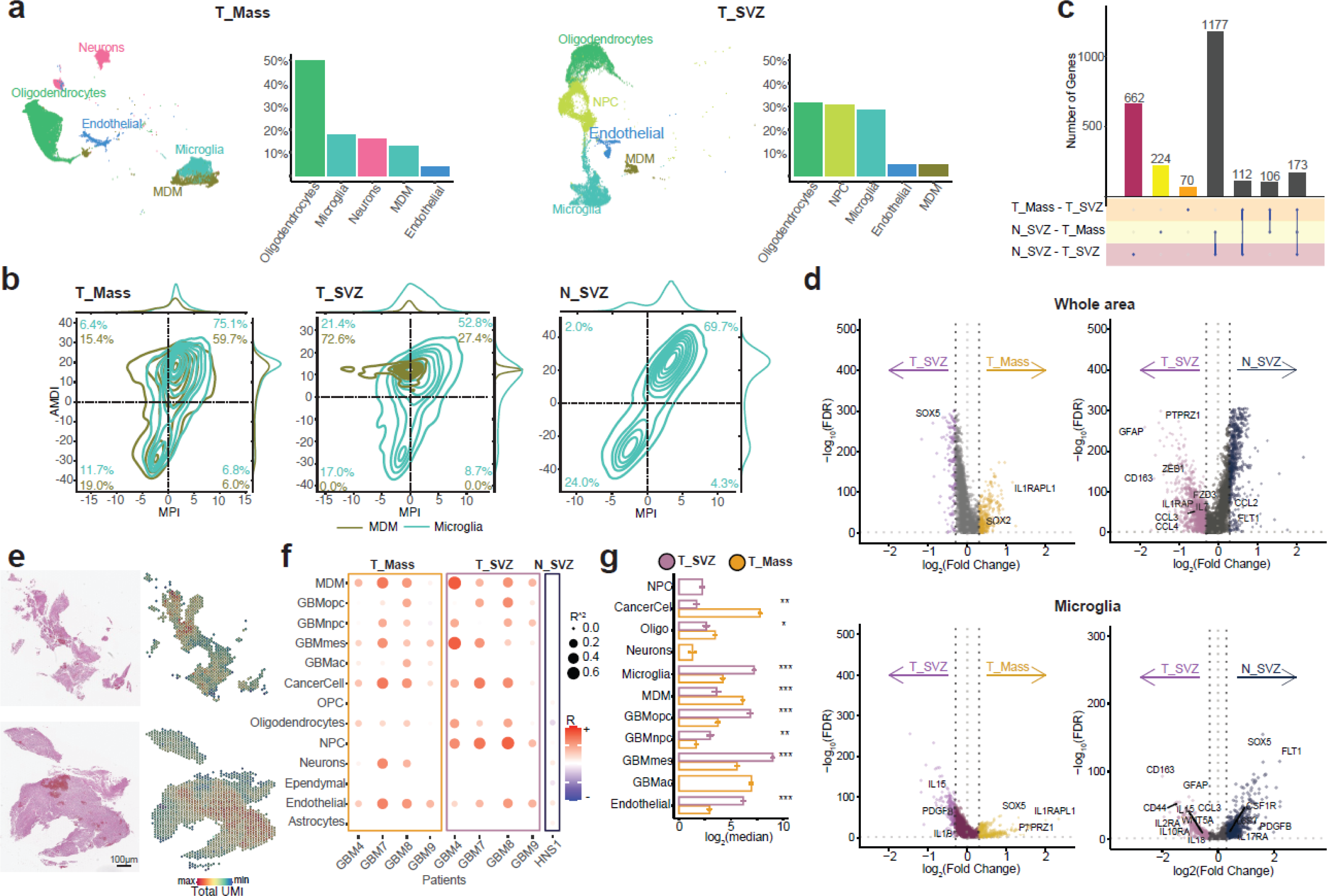
Tumor-supportive microglia are the majority of TAMs in the T_SVZ microenvironment and spatially co-exist with tumor cells. **a,** UMAPs of stromal and immune cell populations in the T_Mass (left) and T_SVZ (right) with proportion of each cell type in the three areas. **b,** MacSpectrum plots of MDM (olive green) and Microglia (green) with percentages calculated for each quadrant in the T_Mass (left), T_SVZ (middle), and N_SVZ (right). AMDI, Activation-induced Macrophage Differentiation Index; MPI, Macrophage Polarization Index. **c,** Upset plot depicting the number of differentially expressed genes in the comparisons among T_Mass, T_SVZ, and N_SVZ. **d,** Volcano plots showing the differentially expressed genes in T_SVZ *versus* T_Mass (left) and T_SVZ *versus* N_SVZ (right) as whole areas, top. Volcano plots showing the differentially expressed genes in microglia only in T_SVZ *versus* T_Mass (left) and T_SVZ *versus* N_SVZ (right), bottom. In all analyses, average log_2_(Fold Change)>0.3 and *p*<0.05 were used. **e,** Representative images of the T_Mass (top) and T_SVZ (bottom) from tissue sections used for spatial transcriptomics. Sections of GBM7 are shown as an example. Images of hematoxylin and eosin-stained tissues (left) and the corresponding digital images (right). Scale bar=100 µm. Color code= minimum to maximum total unique molecular identifier (UMI) for each sample. **f,** Dot plot of Pearson’s spatial correlation between microglia and all other cell types in T_Mass and T_SVZ of 4 GBM patients. Cell types exhibiting spatial correlation with microglia (y-axis). GBM sample areas and normal sample HNS1 (x-axis). **g,** Bar graph showing patterns of spatial dependencies among cell types in the T_Mass (yellow) and in the T_SVZ (purple) from the same GBM patients as in Fig. 3f. All cell types in the two areas were analyzed. Cell types exhibiting spatial dependencies (y-axis) and log_2_ (median) of total spatial dependencies (x-axis). *=*p*<0.05, **=*p*<0.01, ***=*p*<0.001. NPC, neural progenitor cells; OPC, *oligodendrocyte precursor cells;* MDM, monocyte-derived macrophages.

Next, we performed functional characterization of the microglia and MDM in the T_SVZ compared to matched cell populations in the T_Mass. Although M1 is typically synonymous with “proinflammatory/anti-tumor” and M2 with “anti-inflammatory/pro-tumor”, these definitions do not fully represent the functional identity of TAMs in GBM. Thus, we used MacSpectrum^47^ to infer activity of TAMs. Based on input RNA-seq count data, this method determines the Macrophage Polarization Index (MPI) and the Activation Induced Macrophage Differentiation Index (AMDI), both with scores that range from −50 to 50. Instead of a category representation, we mapped macrophage activity onto a biological spectrum using this score-based method. More pro-inflammatory traits are indicated by a higher MPI value, while greater maturity is indicated by a higher AMDI value. Zero was the threshold to designate “pre-activation” or “M0” cells (AMDI < 0, MPI < 0), “M1-transitional” (AMDI < 0, MPI > 0) or “M1-like” cells (AMDI > 0, MPI > 0), and “M2-like” cells (AMDI > 0, MPI < 0), as done by Li *et al.*^47^.

Compared to the T_Mass, microglia of the T_SVZ were more M2-like (from 6.4% to 21.4%) and less M1-like (from 75.1% to 52.8%) and showed an increase in the pre-activation state (from 11.7% to 17.0%, **Fig. 3b left** and **middle**). Similarly, MDM of the T_SVZ were predominantly M2-like compared to those in the T_Mass (from 15.4% to 72.6%) and less M1-like (from 59.7% to 27.4%, **Fig. 3b left** and **middle**). Notably, no MDM exhibited a pre-activation state in the T_SVZ (from 19.0% in the T_Mass to 0.0% in the T_SVZ, **Fig. 3b left** and **middle**). Compared to the N_SVZ, as expected, microglia in this area were predominantly M1-like (69.7%) and in the pre-activation state (24.0%) (**Fig. 3b right**). Overall, microglia in the T_SVZ were prominently tumor-supportive. Thus, TAM-specific mechanisms promoting tumor aggressiveness in the T_SVZ represent novel therapeutic vulnerabilities.

To identify potentially targetable genes in the microglia of the T_SVZ, we conducted gene expression analysis and identified the differentially expressed genes (DEGs) in that area compared with the T_Mass and the N_SVZ. The N_SVZ and the T_SVZ had the highest number of DEGs, while the T_Mass and the T_SVZ had the lowest number of DEGs (**Fig. 3c**). Moreover, the highest number of common DEGs were found between N_SVZ-T_Mass and N_SVZ-T_SVZ (1177 DEGs), while the number of common DEGs between T_Mass-T_SVZ and N_SVZ-T_SVZ, and those between T_Mass-T_SVZ and N_SVZ-T_Mass, were only 112 and 106, respectively. These data suggest that the T_Mass and the T_SVZ express similar genes, while the N_SVZ is characterized by a different gene set (**Fig. 3c**). Notably, among the DEGs, the CSC master regulator *SOX2*^41,48^ was significantly downregulated in the N_SVZ *versus* T_Mass and *ZEB1* was significantly downregulated in the N_SVZ *versus* T_SVZ, thus supporting the results using SCENIC (**Fig. 2d left**). Moreover, TFs involved in promotion of neuronal differentiation^49^ showed two distinct patterns: while *SOX4* and *SOX11* were overexpressed in the T_Mass *versus* N_SVZ and in the T_SVZ *versus* N_SVZ, *SOX5* was significantly upregulated in the N_SVZ *versus* T_Mass comparison (**Suppl. Table 4**). Of note, *SOX2*, *SOX4*, and *SOX11* were upregulated in the T_Mass *versus* T_SVZ, and *SOX5* was downregulated in the T_Mass *versus* T_SVZ. Of note, *COL1A1* (a central gene in the dynamic organization of glioma mesenchymal transformation^50^) was downregulated in the T_Mass *versus* T_SVZ, further supporting our observation that the T_SVZ microenvironment is characterized by a mesenchymal signature.

When considering overexpressed genes in the T_SVZ as a whole area (*i.e.* all identified cell types in this microenvironment), *SOX2* was upregulated in the T_Mass *versus* T_SVZ (**Fig. 3d top left**). The gene encoding for the IL-1R accessory protein (IL-1RAcP) was significantly upregulated in the T_SVZ *versus* N_SVZ (**Fig. 3d top right**). Moreover, in addition to *GFAP*, *CD163*, *PTPRZ1*, and *ZEB1*, we also observed that *FZD3*, encoding a seven-transmembrane domain receptor of the ncWNT pathway, was significantly upregulated in the T_SVZ *versus* N_SVZ and in the T_Mass *versus* N_SVZ (**Fig. 3d top right** and **Suppl. Fig. 7 left**).

We then performed the same analysis on the microglia only (**Suppl. Table 5**). The inflammatory cytokine *IL1B*^51–58^ was significantly upregulated in the T_SVZ *versus* T_Mass, suggesting that microglia in the T_SVZ are more inflammatory than those in the T_Mass (**Fig. 3d bottom left**). Two other inflammatory cytokines were upregulated in the T_SVZ microglia: *IL15* was significantly upregulated in the T_SVZ *versus* T_Mass and in the T_SVZ *versus* N_SVZ (**Fig. 3d bottom left** and **right**), and *IL18* was significantly upregulated in the T_SVZ *versus* N_SVZ (**Fig. 3d bottom right**), thus confirming that T_SVZ microglia are prominently tumor-supportive and inflammatory, consistent with the results of MacSpectrum (**Fig. 3b**). We confirmed these results by analyzing gene expression of markers of ‘homeostatic’, ‘activated’, and ‘inflammatory’ microglia (**Suppl. Fig. 8**). In addition to *CD163*, *CD44*, *GFAP*, *IL2RA*, and other genes, the pro-migration and pro-invasion non-canonical WNT (ncWNT) ligand *WNT5A*^59–63^ was significantly upregulated in the T_SVZ *versus* N_SVZ and in the T_Mass *versus* N_SVZ microglia only comparisons (**Fig. 3d bottom right** and **Suppl. Fig. 7 right**). Of note, high expression of *WNT5A* in glioma has been correlated with increased presence of TAMs^64^. Moreover, the putative receptor of *WNT5A* is *FZD3*, which was significantly upregulated in the T_SVZ *versus* N_SVZ and in the T_Mass *versus* N_SVZ whole area comparisons (**Fig. 3d top right** and **Suppl. Fig. 7 left**).

These results led us to explore the spatial distribution of microglia and tumor cells in cellular neighborhoods of the T_SVZ and the T_Mass microenvironments. We performed spatial transcriptomics using samples from 4 GBM patients in our cohort with large enough T_SVZ and T_Mass tissues for analyses (**Fig. 3e**). We also profiled the HNS1 sample (one of our two N_SVZ samples). By InferCNV, cell type annotation, and Cell2Location, we first confirmed that most of the cells in the T_Mass and the T_SVZ were tumor cells (except for the GBM4 samples) and that the HNS1 contained normal cells only (**Suppl. Fig. 9**). As expected, we found a higher percentage of microglia and GBMmes cells in the T_SVZ samples compared to corresponding T_Mass samples (**Suppl. Fig. 10**). We then quantified the spatial correlations for each patient and each microenvironment; microglia exhibited a strong spatial correlation with tumor cells except for GBM9 (**Fig. 3f**). We also observed different patterns of correlations with tumor and normal cells between the T_Mass and the T_SVZ of each patient (**Suppl. Fig. 11**). Microglia established a stronger spatial correlation with MDM in the T_SVZ than in the corresponding T_Mass, and a spatial correlation with NPCs was observed in all T_SVZ samples (**Fig. 3f**). Of note, correlation analyses of microglia in the HNS1 sample revealed significantly weaker correlations with all cell types (**Fig. 3f**). When we examined patterns of spatial dependencies among cell types^65^, we observed that microglia in the T_SVZ had a high likelihood of communication with tumor cells and were both sender and receiver cells in 2 of 4 T_SVZ samples. By contrast, in the corresponding T_Mass samples, microglia were only receiver cells, similar to the HNS1 sample (**Suppl. Fig. 12**).

Altogether, these data indicate that the T_SVZ microenvironment is characterized by tumor-supportive microglia that secrete inflammatory cytokines, such as IL-1β, and the pro-migration and pro-invasion ncWNT ligand Wnt-5a. Microglia in the T_SVZ established spatial dependencies predominantly with tumor cells.

### Microglia, not MDM, establish cell-to-cell interactions with tumor cells in the T_SVZ microenvironment and are predicted to express IL-1β and Wnt-5a

The spatial co-existence of tumor-supportive, inflammatory microglia and tumor cells in the T_SVZ microenvironment prompted us to further examine their interactions. First, we annotated each identified cluster in the T_Mass, T_SVZ, and N_SVZ to identify interactions at high resolution (**Fig. 4a top**). Next, we examined the total number of inferred interactions. The N_SVZ had the lowest number of interactions (221), the T_SVZ had an intermediate number (326), and the T_Mass had the highest number (653) (**Suppl. Fig. 13**). These findings correlated with the number of identified clusters (**Fig. 4a top**) and suggest that more heterotypic cellular microenvironments, such as those of the T_Mass and the T_SVZ (**Fig. 1f bottom**), contribute to increased cell-to-cell interactions. We then analyzed the number of the incoming and outgoing interactions among cell types. While MDM exhibited a cell-to-cell communication network in the T_Mass (**Fig. 4a bottom left**), they establish only a few, weak interactions in the T_SVZ (**Fig. 4a bottom middle**). In contrast, microglia established cell-to-cell communication networks in both the T_Mass (**Fig. 4a bottom left**) and the T_SVZ (**Fig. 4a bottom middle**). In the T_SVZ, microglia showed interactions with different cell types, including tumor cells of the GBMmes state (**Fig. 4a bottom middle**). These data on microglia in the T_SVZ seem to reflect the ability of microglia in the N_SVZ to be highly interactive within the microenvironment and establish a complex cell-to-cell communication network with many cell types (**Fig. 4a bottom right**).

**Figure 4.**
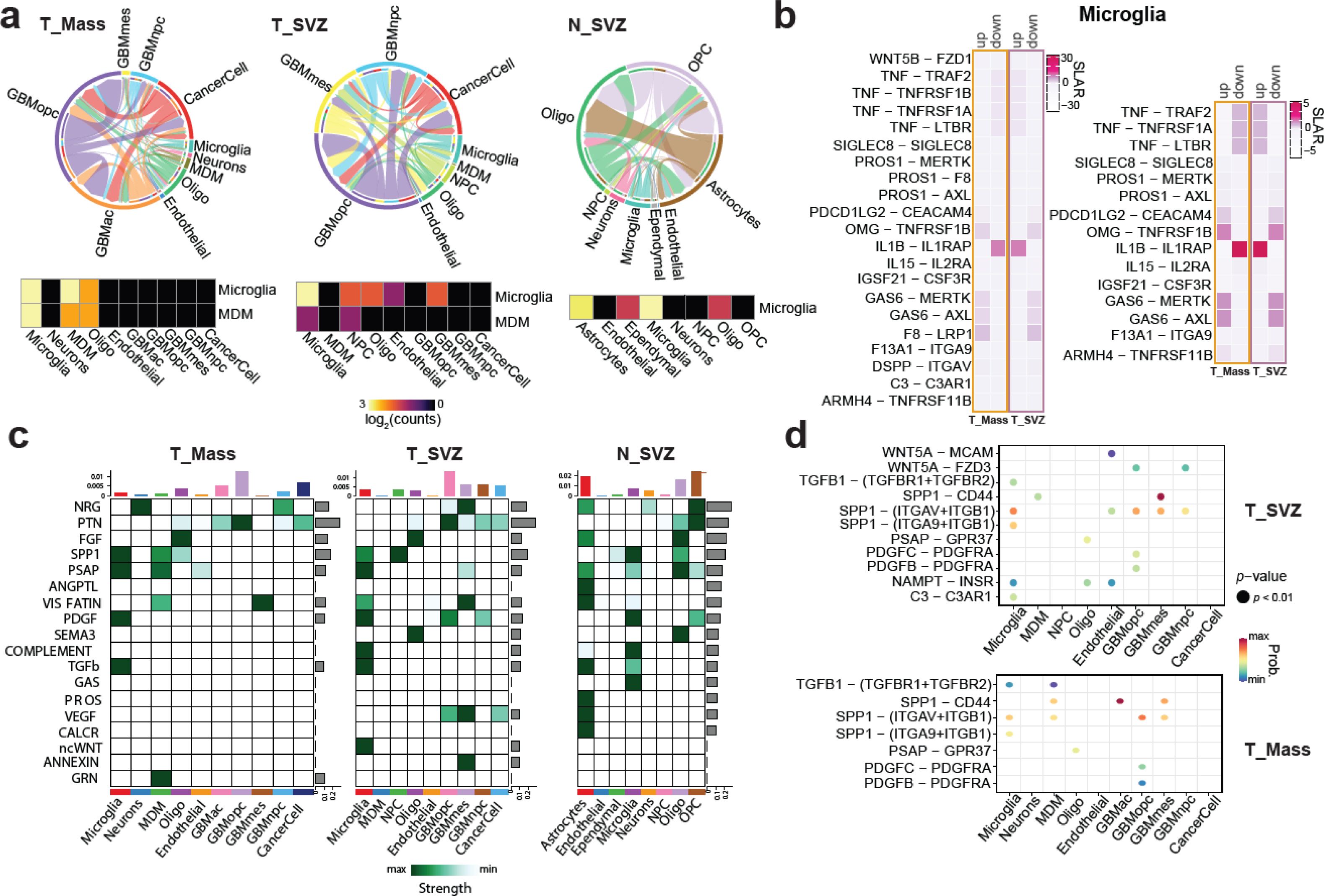
Microglia, not MDM, establish cell-to-cell interactions with tumor cells in the T_SVZ microenvironment and are predicted to express IL-1β and Wnt-5a. **a,** Incoming and outgoing interactions in the cell-to-cell communication network. Number of interactions between any two cell types shown as circle plots for the T_Mass (top left), T_SVZ (top middle), and N_SVZ (top right). Heatmaps showing the cell-to-cell interactions of microglia and MDM with the other cell types and with themselves for the T_Mass (bottom left), T_SVZ (bottom middle), and N_SVZ (bottom right). No MDM were present in the N_SVZ, hence only the heatmap for microglia is shown. Color code indicates the log_2_ (counts) between 0 and 3. **b,** Heatmaps showing the predicted interactions between sender (left) and receiver (right) cells in the microglia cluster of the T_SVZ. Color code indicates the scaled ligand activity in receiver (SLAR) cells. **c,** Heatmaps of outgoing signaling pathways in the T_Mass (left), T_SVZ (middle), and N_SVZ (right). Color code indicates minimum to maximum strength of each signaling pathway. **d,** Dot plots of ligand-receptor prediction analysis between microglia as ligand-expressing cells and any other cell types (x-axis) in the T_SVZ (top) and in the T_Mass (bottom). Color code=minimum to maximum probability. NPC, neural progenitor cells; OPC, *oligodendrocyte precursor cells;* MDM, monocyte-derived macrophages.

We next studied predicted interactions involving microglia of the T_SVZ. First, we examined predicted interactions within this cell type (microglia to microglia). Activity of the IL-1β –IL-1RAcP pathway was strongly upregulated in ‘sender’ (**Fig. 4b left**) and ‘receiver’ (**Fig. 4b right**) cells of the microglia in the T_SVZ compared to microglia of the T_Mass. Second, we examined all predicted incoming (**Suppl. Fig. 14**) and outgoing signaling patterns in the T_Mass (**Fig. 4c left**), the T_SVZ (**Fig. 4c middle**), and the N_SVZ (**Fig. 4c right**). Among the incoming signaling pathways of microglia in the T_SVZ, Sema3 and Annexin had the highest relative strength and were specific to this area compared to the T_Mass and the N_SVZ (**Suppl. Fig. 14**). Consistent with the DEG analysis (**Fig. 3d bottom**), among the outgoing signaling pathways of microglia in the T_SVZ, ncWNT exhibited the highest relative strength and was specific to this area compared to the T_Mass and the N_SVZ (**Fig. 4c middle**). Of note, ncWNT was also an incoming signaling pathway specific of T_SVZ and exhibited the highest relative strength in clusters of the GBMnpc and GBMopc states (**Suppl. Fig. 14**), suggesting a ncWNT-mediated interaction between microglia and tumor cells in the T_SVZ.

We then performed ligand-receptor prediction analyses between microglia and any other cell type of the T_SVZ. *WNT5A* is a predicted ligand of microglia, and its predicted receptors are *MCAM* and *FZD3* expressed by endothelial cells and tumor cells of the GBMopc and GBMnpc states, respectively (**Fig. 4d top**). These *WNT5A*-centered predictions were specific to the T_SVZ and absent in the T_Mass (**Fig. 4d bottom** and **Suppl. Fig. 15**), in agreement with the incoming and outgoing signaling analyses in the T_SVZ (**Fig. 4c middle** and **Suppl. Fig. 14**). The *SPP1*-*CD44* ligand-receptor combination had the highest communication probability between microglia and tumor cells of the GBMmes state in the T_SVZ (**Fig. 4d top**) and between microglia/MDM and tumor cells of the GBMac and the GBMmes states in the T_Mass (**Fig. 4d bottom** and **Suppl. Fig. 15**). This is consistent with published work showing that: (i) *SPP1*-*CD44* signaling is present in the glioma perivascular niche^66^, (ii) *SPP1* is upregulated^67^ and secreted by TAMs in glioma^68^, and (ii) *SPP1*-*CD44* signaling is between TAMs and glioma cells^69^, specifically with GBMmes tumor cells^70^.

Overall, our results revealed that microglia in the T_ SVZ establish cell-to-cell interactions within their cell population and with tumor cells and identify microglia-specific pathways of communications.

### IL-1β/IL-1RAcP and Wnt-5a/Frizzled-3 are therapeutic targets in the T_SVZ microenvironment

Based on our results above revealing that *IL1B* and *WNT5A* are significantly upregulated in the T_SVZ microglia *versus* T_Mass and N_SVZ, respectively (**Fig. 3d bottom left and right**) and that their predicted receptors *IL1RAP* (**Fig. 4b**) and *FZD3* (**Fig. 4d top**) are also significantly upregulated in the T_SVZ *versus* N_SVZ comparison as whole areas (**Fig. 3d top right**), we next performed functional studies of the IL-1β /IL-1RAcP and Wnt-5a/Frizzled-3 pathways. Given the recognized inflammatory and tumor-supportive role of IL-1β^51–58^, and the pro-migration and pro-invasion functions of Wnt-5a^59–63^, we surmised that strategies to target these two pathways could reveal therapeutic vulnerabilities in the T_SVZ microenvironment.

First, we defined the cell type expression and co-expression levels of *IL1B* and *WNT5A* and of *IL1RAP* and *FZD3* in the T_SVZ (**Fig. 5a**). While *WNT5A* and *IL1B* were almost exclusively expressed by microglia (**Fig. 5b**), *FZD3* was predominantly expressed by tumor cells of the GBMnpc and GBMopc states, and *IL1RAP* was predominantly expressed by tumor cells of those two states and microglia (**Fig. 5b**). The cell type expression of *WNT5A, IL1B*, *FZD3*, and *IL1RAP* in the T_SVZ overlapped only partially with the expression of the same genes in the T_Mass and the N_SVZ (**Suppl. Fig. 16**).

**Figure 5.**
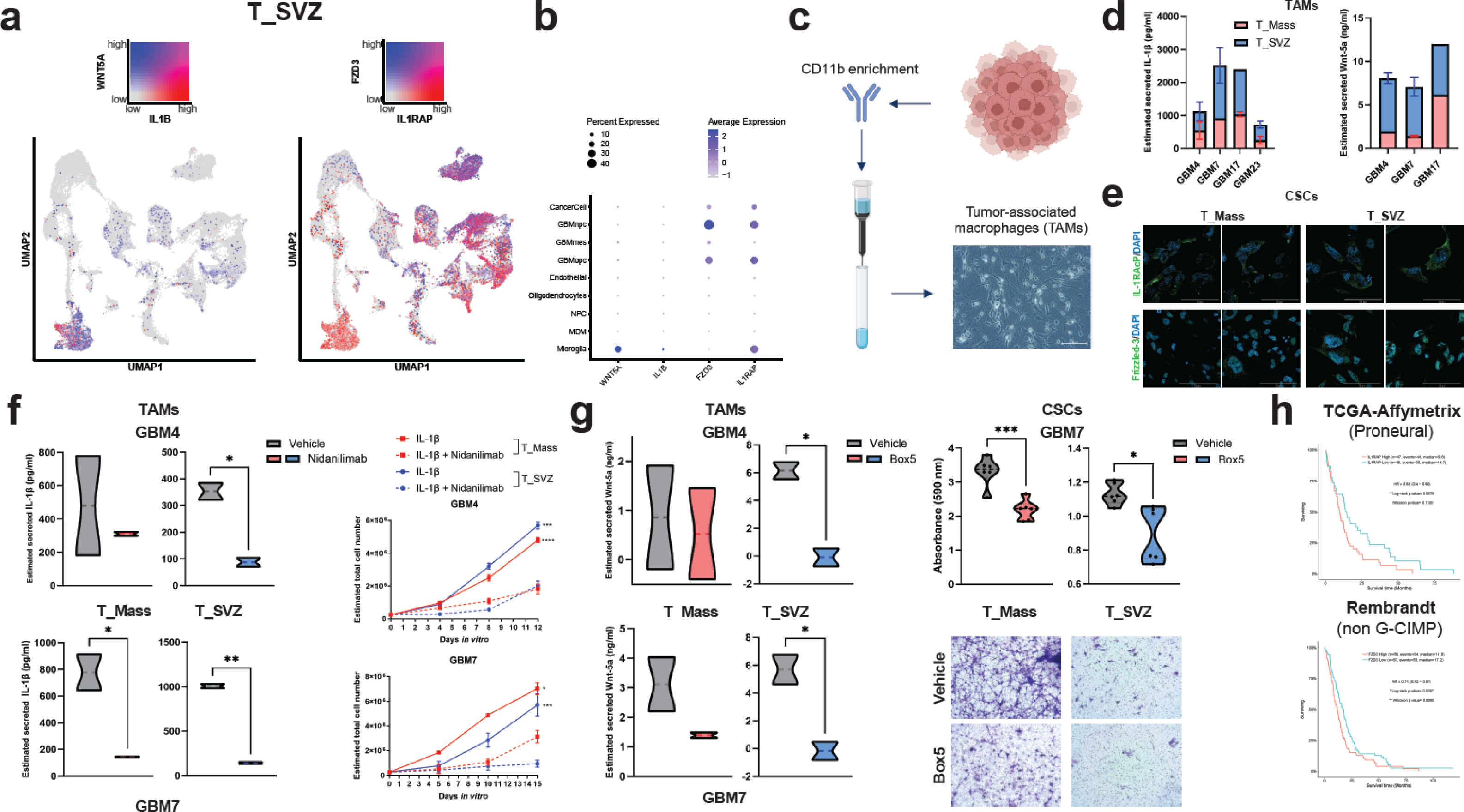
IL-1β/IL-1RAcP and Wnt-5a/Frizzled-3 are therapeutic targets in the T_SVZ microenvironment. **a,** UMAPs of the T_SVZ showing expressions of *IL1B*, *WNT5A* (left) and *IL1RAP* and *FZD3* (right) and gene co-expression (top). Cell type annotation of these UMAPs is shown in Fig.1f. **b,** Dot plot showing cell type expression of *WNT5A*, *IL1B*, *FZD3*, and *IL1RAP* in the T_SVZ. **c,** Schematic of the cd11b enrichment strategy for isolation of tumor-associated macrophages (TAMs). Created with BioRender.com. **d,** Bar graph of estimated IL-1β (left) and Wnt5-a (right) secreted from the TAMs isolated from the T_Mass and the T_SVZ of GBM4, 7, 17, and 23 (the latter only for IL-1β; mean±s.e.m.). **e,** Representative images of CSCs isolated from GBM4 and 7 and stained for IL1-RAcP (top) and Frizzled-3 (bottom). Counterstained with DAPI. Scale bars=50 µm. **f,** Truncated violin plots of estimated secreted IL-1β by TAMs isolated from T_Mass and T_SVZ of GBM4 (top) and GBM7 (bottom) after Nidanilimab treatment. *=*p*<0.05, **=*p*<0.01 (left). Growth curve analysis (mean±s.e.m.) of CSCs isolated from T_Mass and T_SVZ of GBM4 (top) and GBM7 (bottom) and treated with IL-1β and IL-1β+Nidanilimab (right). Non-linear regression analysis was performed to assess the proliferative potential. *=*p*<0.05, ***=*p*<0.001, ****=*p*<0.0001 IL-1β *versus* IL-1β+Nidanilimab. **g,** Truncated violin plots of the estimated secreted Wnt-5a by TAMs isolated from T_Mass and T_SVZ of GBM4, and 7 after treatment with Box5. *=*p*<0.05 (left). Quantification of cell migration based on absorbance of eluted crystal violet used in transwell assays of CSCs isolated from T_Mass and T_SVZ of GBM7 and exposed to conditioned medium of matched TAMs treated with Box5. *=*p*<0.05, ***=*p*<0.001 (top right). Representative images of migrated CSCs in the vehicle and Box5-treated conditions and stained with crystal violet. Stained spots are pores on the membranes (bottom right). **h,** Correlation between *IL1RAP* (top) and *FZD3* (bottom) expression levels and patient survival in the TCGA (Affymetrix platform, proneural subtype) and Rembrandt datasets (non G-CIMP status), respectively. The survival analysis was performed using the GlioVis data portal^80^.

Second, similar to previous analyses of TAM expression in human gliomas, we isolated TAMs using the cd11b enrichment strategy based on immunomagnetic microbeads decorated with recombinantly engineered antibody fragments (**Fig. 5c**) and quantified levels of IL-1β (**Fig. 5d left**) and Wnt-5a (**Fig. 5d right**) secreted by TAMs of matched T_Mass and T_SVZ of 4 GBM patients. Using immunofluorescence staining, we confirmed that IL-1RAcP and Frizzled-3 were expressed by CSCs (**Fig. 5e**). For IL-1RAcP, we also confirmed expression at the level of TAMs (**Suppl. Fig. 17**). We observed higher expression of IL-1RAcP in TAMs isolated from the T_SVZ compared to matched TAMs of the T_Mass (**Suppl. Fig. 17**) and of Frizzled-3 in the CSCs isolated from the T_SVZ compared to matched CSCs of the T_Mass (**Fig. 5e**).

Third, we tested the effects of IL-1β/IL-1RAcP and Wnt-5a/Frizzled-3 inhibition *in vitro*. To evaluate the impact of IL-1β/IL-1RAcP inhibition in TAMs, isolated cd11b-enriched cells from matched T_SVZ and the T_Mass of 2 GBM patients were treated for 48 hours with the anti-IL-1RAcP fully humanized monoclonal antibody Nidanilimab (currently being tested in multiple clinical trials^71–73^). We observed significantly reduced IL-1β secretion in Nidanilimab-treated TAMs of the T_SVZ in GBM4 (**Fig. 5f top**) and in both the T_Mass and the T_SVZ for GBM 7 (**Fig. 5f bottom**). Next, we evaluated the impact of IL-1β/IL-1RAcP inhibition on CSCs. Proliferation of CSCs isolated from the T_Mass and the T_SVZ was significantly reduced after treatment with IL-1β and Nidanilimab (**Fig. 5f right**) but not with Nidanilimab alone (data not shown), with a more pronounced effect in the T_SVZ compared to T_Mass (**Fig. 5f right**). These data suggest that mimicking the secretion of IL-1β by TAMs is critical for IL-1β/IL-1RAcP inhibition. To test the impact of Wnt-5a/Frizzled-3 inhibition, we performed *in vitro* transwell experiments using CSCs isolated from the T_Mass and the T_SVZ of 3 GBM patients. Initially, TAMs isolated from matched T_SVZ and the T_Mass of the 3 patients were treated for 48 hours with Box5, a Wnt-5a antagonist. Notably, secretion of Wnt-5a was significantly reduced upon Box5 treatment of TAMs isolated from the T_SVZ (**Fig. 5g left**). Exposure of CSCs to conditioned medium of treated or untreated TAMs with Box5 significantly reduced CSC invasion through the transwell (**Fig. 5g right** and **Suppl. Fig. 18**).

To evaluate the clinical significance of our findings, we used The Cancer Genome Atlas (TCGA) and the Rembrandt GBM datasets to perform analysis of overall survival based on the expression levels of *IL1RAP* and *FZD3*. High expression of *IL1RAP* and *FZD3* were associated with shorter survival in the TCGA samples (Affymetrix platform, proneural subtype), and in the Rembrandt, non G-CIMP status datasets, respectively (**Fig. 5h**).

Collectively, these results indicate that the IL-1β/IL-1RAcP and Wnt-5a/Frizzled-3 represent therapeutic targets in the T_SVZ and have prognostic relevance in subsets of GBM patients.

## Discussion

We built a sn-RNAseq-based microenvironment landscape of the T_SVZ using samples from 15 GBM patients. We comprehensively compared T_Mass samples isolated from the same patients and used two histologically normal SVZ samples as controls. We identified a *ZEB1*^40–42^-based mesenchymal signature in the T_SVZ and a tumor-supportive microglia population, which represent the vast majority of TAMs in the T_SVZ microenvironment. These cells spatially co-exist and establish cell-to-cell interactions with tumor cells. We systematically characterized these interactions both *in silico* and *in vitro* and identified two pathways, IL-1β/IL-1RAcP and Wnt-5a/Frizzled-3, representing novel targets in the T_SVZ microenvironment.

Collectively, our findings indicate that the SVZ represents a distinct microenvironment in GBM patients, which is deprived of the GBMac state, is characterized by a mesenchymal signature, and is enriched in tumor-supportive and inflammatory microglia. To the best of our knowledge, this study is the first to identify potential therapeutic targets in the T_SVZ of GBM patients.

While IL-1β is among the most-well characterized inflammatory cytokines in GBM, TAMs secreting IL-1β support tumor growth^51–58^, and microglia (among other cell types) are enriched in an IL-1β inflammatory program^74^, it is still unknown whether TAMs secrete Wnt-5a and if a Wnt-5a-mediated crosstalk exists between TAMs and tumor cells. However, in addition to the known pro-migration and pro-invasion functions of Wnt-5a^59–63^, it has been reported that the expression of Wnt-5a in human glioma is positively correlated with the presence of TAMs^64^.

Our results show that *in vitro* targeting of the IL-1β/IL-1RAcP and Wnt-5a/Frizzled-3 pathways significantly reduced the ability of CSCs of the T_SVZ to proliferate and migrate. Of note, inhibition of the IL-1β/IL-1RAcP pathway with an anti-IL-1RAcP fully-humanized monoclonal antibody, Nidanilimab, is being evaluated in multiple clinical trials of colorectal cancer, non-small cell lung cancer, pancreatic cancer, triple-negative breast cancer^71–73^, and biliary tract cancer (https://cantargia.com/en/press-releases/cantargia-reports-treatment-of-first-triple-negative-breast-cancer-patient-in-trifour-study).

Given the limited efficacy of current treatments for GBM patients, our results provides evidence of novel potential therapeutic opportunities to target the T_SVZ microenvironment. Previous work identified associations between GBMmes and MDM, and between GBMac and microglia^75^, as well as a more abundant specific MDM population in GBM with radiographic contact to the ventricle, whereas microglia are enriched in GBM showing no ventricular contact^76^. However, none of these studies was performed on tissues sampled directly from the SVZ of GBM patients.

Although microglia undergo changes in gene expression from ‘homeostatic’ to ‘activated’ in CNS diseases, their diversity and functional roles in human GBM are not fully understood^77,78^. By distinguishing between MDM and microglia, elegant works on the brain tumor microenvironment started to uncover the phenotype of microglia in IDH wild-type/mutant gliomas, and in brain metastases^20,21^ and suggested that microglia exhibit an ‘activated’ phenotype in GBM. In addition, two independent groups have shown that in GBM patients, subsets of microglia upregulate inflammatory (including *IL1B*) and proliferative genes^56^ and are characterized by *VEGF*- and *CD163*-expressing cells^79^, suggesting a tumor-supportive function whose mechanisms are still unknown.

Our work has some limitations: we have not analyzed all of the cellular cross-talks mediated by IL-1β, and we have not evaluated the therapeutic efficacy of inhibiting the IL-1β/IL-1RAcP or the ncWNT (Wnt-5a/Frizzled-3) pathways *in vivo*. While inhibition of secreted factors can be challenging due to the dynamics of secretion and the cross-talks between different cell types, disruption of cellular interactions at the level of receptors through blocking antibodies could be a more successful approach. Exploration of this and other therapeutic opportunities based on T_SVZ-specific targets could lead to more effective therapies for GBM patients. Of note, our patient cohort was not powered for the analysis of sex-related differences. Future work will evaluate the therapeutic efficacy of inhibiting the IL-1β/IL-1RAcP and the ncWNT (Wnt-5a/Frizzled-3) pathways in a sex-specific manner.

## Data availability

Raw and processed data to support the findings of this study have been deposited in GEO and will remain private until publication. All processed data, UMAP coordinates, and annotations are available to download and inspect at the BioCM GitHub portal through a shiny app upon reasonable request to the corresponding author.

## Acknowledgments

This work was supported by The Ben and Catherine Ivy Foundation Translational Adult Glioma Award and The Robert M. Faxon Jr. Endowed Professorship in Neuro-Oncology to S.G.M.P. Y.L.-M. is partially supported by NIH grant P20GM121176. V.A. was supported by the PREP program of the University of New Mexico. S.S., B.G., and S.B. are supported by the CNDD Genomics and Bioinformatics Core at MUSC (NIH grant P20GM148302). The authors are grateful to the Chavez family for the postmortem brain donation.

This work was also partially supported by UNM Comprehensive Cancer Center Support Grant NCI P30CA118100 and made use of the Analytical and Translational Genomics Shared Resource, which receives additional support from the State of New Mexico, and the Human Tissue Repository and Tissue Analysis Shared Resource, also funded by the Department of Pathology of the University of New Mexico. The authors also acknowledge the use of the UNM Autophagy, Metabolism, and Inflammation core funded by NIH grant P20GM121176 and are grateful to the UNM Hospital Neurosurgery Research Coordinators and Managers for assistance with the patient sample collection, to Sharina Desai for helping with the resources of the Autophagy, Metabolism, and Inflammation core, and to Kel Cook and Cathleen Martinez for the help with the set-up of the first spatial transcriptomic test.

The authors also acknowledge Singulomics and 3D Genomics for the single-nuclei RNA sequencing and spatial transcriptomics data generation and are grateful to Karen Klein for valuable help with the editing of this manuscript.

## Author contributions

S.G.M.P. designed and supervised the study. S.G.M.P, Y.L.-M., S.S., B.G., and S.B. wrote the manuscript. S.G.M.P. coordinated experiments. Y.L.-M., V.A., and F.M. performed the experiments and contributed to data interpretation. S.V. assisted with the collection and curation of clinical data. S.S., B.G., D.M., and S.B. wrote all original code used in the study and performed bioinformatic analysis. E.P., M.K., M.O.C., and C.A.B. provided patient tissue samples. All authors reviewed and approved the manuscript.

## Materials and Methods

### Glioblastoma sample collection

Patient informed consent was obtained through the Neurosurgery Clinics at the University of New Mexico Hospitals and at the University of Mississippi Medical Center. Tissue collection protocols were IRB-approved. Patient clinical and molecular information is provided as Supplementary Information.

### Genomic analysis of the glioblastoma samples

Genomic DNA extraction from glioblastoma tissues was performed using the DNeasy Blood & Tissue Kit (Qiagen, Cat. No. 69506). To perform targeted gene sequencing the AmpliSeq Cancer Hotspot Panel v2 (Thermo Fisher Scientific, Cat. No. 4475346), which covers approximately 2,800 COSMIC mutations in 50 genes, was used. Libraries were prepared using Ion AmpliSeq Library Kit 2.0 (Thermo Fisher Scientific, Cat. No. 4480441) and Ion XPress Barcode Adapters (Thermo Fisher Scientific, Cat. No. 4471250) following the manufacturer’s instructions. GBM samples were amplified using 10 ng of input DNA. Libraries were purified and size-selected using Agencourt AMPure XP beads (Beckman Coulter, Cat No. A63880), quantified using the Qubit 2.0 Fluorometer (Thermo Fisher Scientific, Cat. No. Q32866) and Agilent 2100 Bioanalyzer (Agilent, Cat. No. G2939BA), and diluted to 50 pM. Diluted libraries were loaded onto Ion 550 chips using an Ion Chef instrument and sequenced on the Ion S5 XL (Thermo Fisher Scientific).

The VCF files from the Ion Torrent Variant Caller were annotated using SNP EFF v4.3.1t^1^. Mutations with allele frequencies below 0.05 were filtered out, as well as nonsense mutations and those with known non-pathogenic variants. For analysis of the genetic drivers for each sample, any mutations for genes *CDKN2A*, *PDGFRA*, and *TP53* were plotted in a heatmap along with genetic and copy number data for the same genes along with *EGFR* and *NF1.* The genetic data for each gene is derived from the single-cell RNA expression data, then averaged for each sample, and Z-scored. The copy number data were obtained from InferCNV^2^ and similarly averaged for each sample and Z-scored.

### Single-nucleus RNA sequencing

Single-nucleus RNA sequencing and analysis were conducted by Singulomics Corporation (https://singulomics.com/, Bronx NY). In summary, frozen human tissue samples were homogenized and lysed with Triton X-100 in RNase-free water for nuclei isolation. The isolated nuclei were purified, centrifuged, and resuspended in PBS with BSA and RNase Inhibitor. The nuclei were diluted to 700 nuclei/ul and loaded to 10x Genomics Chromium Controller to encapsulate single nuclei into droplet emulsions following the manufacturer’s recommendations (Pleasanton, CA, United States). Library preparation was performed according to the instructions in the Chromium Next GEM 3’ Single Cell Reagent kit v3.1. Amplified cDNAs and the libraries were measured by Qubit dsDNA HS assay (Thermo Fisher Scientific, Wilmington, DE) and quality was assessed by BioAnalyzer (Agilent Technologies, Santa Clara, CA). Libraries were sequenced on a NovaSeq 6000 instrument (Illumina, San Diego, CA, United States), and reads were subsequently processed using 10x Genomics Cell Ranger analytical pipeline and human GRCh38 reference genome with introns included in the analysis. Dataset aggregation was performed using the cellranger aggr function normalizing for the total number of confidently mapped reads across libraries. Specifically, raw base call (BCL) files were analyzed using CellRanger (v7.0.0)^3^. The “mkfastq” command was used to generate FASTQ files and the “count” command was used to generate raw gene-cell expression matrices. Ambient RNA contamination was inferred and removed using CellBender (v0.2.0) with standard parameters. Human genome hg38 was used for the alignment and gencode.v42 gtf file was used for gene annotation and coordinates^4^. Data from the three areas (N_SVZ, T_SVZ, and T_Mass) were analyzed individually and subsequently integrated. Samples from patients were combined in R using the Read10X function from Seurat package(v4.3.0)^5^, and an integrated Seurat object was generated. Filtering was conducted by retaining cells that had unique molecular identifiers (UMIs) less than 25000 and had mitochondrial content less than 5 percent. Doublets were removed using *scDblFinder* (v1.12.0)^6^. To account for biological and technical batch differences between patients, we used *SCTransform*. This approach was used for count normalization, initial integration, and to identify highly variable genes^7^. We further removed batch effect between the single-cell transcriptome expression matrices of the filtered high-quality cells using Harmony (v0.1.1)^8^. 3000 variable genes were selected for principal components analysis (PCA). The top 30 significant principal components (PCs) and a resolution of 0.3 for Louvain clustering were selected for Uniform Manifold Approximation and Projection (UMAP) and visualization of gene expression. Cluster marker genes were identified using FindAllMarkers using the Wilcoxon Rank Sum test with the standard parameters. N_SVZ is from two patients HNS1 and HNS2 with a total of 19988 protein coding genes and 8772 cells. T_SVZ is from 12 patients GBM7B, GBM8B, GBM9B, GBM10B, GBM12B, GBM16B, GBM17B, GBM20B, GBM22B, MIS1B, MIS2B, MIS3B with a total of 19988 genes and 37360 cells. T_Mass is from 14 patients GBM7A, GBM8A, GBM9A, GBM10A, GBM12A, GBM14A, GBM16A, GBM17A, GBM18A, GBM20A, GBM22A, MIS1A, MIS2A, MIS3A with a total of 19988 protein-coding genes and 67801 cells. For N_SVZ Cell annotation was performed using two different approaches: 1) scType, an ultrafast unsupervised method for cell type annotations^9^, and 2) Manual curation by gene markers to reflect the prediction results. The three areas were subsequently integrated with Harmony (v0.1.1)^8^ using 30 PCA and 3000 most variable genes.

### Definition of malignant cells

Putative malignant cells were identified using InferCNV analysis^2^ with the following parameters - denoise TRUE, default Hidden Markov Model (HMM) settings, and a value of 0.1 for “cutoff” and the N_SVZ clusters were used are reference. Each CNV was annotated to be either a gain or a loss. Tumor cell clusters were classified based on the cell states by Neftel *et al.*^10^ using their meta module gene markers, SCpubr (V2.0.1) was used to visualize the resultant enrichment. The clusters that showed an enrichment for a cell state were labeled as GBMopc (OPC-like), GBMac (AC-like), GBMmes (MES-like), GBMnpc (NPC-like), other clusters that either showed enrichment for two or more cell states or no enrichment for any cell state were labeled as GBMcc (CancerCell). Healthy cells were manually curated by using canonical gene markers: Astrocytes - “GFAP”,“AQP4”, Microglia - “MEF2C”,“P2RY12”, Neurons - “SYNPR”,“CNR1”, “SYT1”, Oligodendrocytes - “MOG”,“MBP”, OPC - “VCAN”,“SOX5”, Endothelial - “VWF”,“ABCB1”, Ependymal - “SPARCL1”,“S100B”.

### Cellular dynamics

Cellular dynamics were analyzed using Cellrank v2.0.2^11^ and CellRank v1^12^. Each region’s dataset was preprocessed using the RNA Velocity steps along with gene imputation using MAGIC^13^. For each area (T_Mass and T_SVZ), a Velocity Kernel was computed separately for tumor and normal cells using all genes in each dataset. A GPCCA^14^ estimator was used to compute macrostates, including the initial and terminal ones for each subset. The number of macrostates was determined using the Schurr decomposition within CellRank individually, and 6 states were selected for each subset. Fate probabilities were computed using the ‘direct’ solver with ‘use_petsc’ set to True. The ‘ilu’ preconditioner was also used in this step. To create heatmaps, a GAM model with 6 knots was used, and the genes selected for the heatmap correspond to the transcription factors with the top 10 regulon specificity scores per cell class.

### Gene regulatory networks

Transcription factor regulatory networks were computed using pySCENIC (v0.12.1)^15^. Each of the T_Mass and the T_SVZ were divided into sections containing only tumor and normal cells respectively, and each of these four datasets was processed separately. The gene regulatory networks were computed using the grn algorithm for each subsample. The ctx method was then run to find enriched motifs using the hg38_10kbp up/down and hg38_500bp up/down motif feather databases downloaded from https://resources.aertslab.org/cistarget/. Last, the aucell method was run to calculate regulome enrichment for each cell in each dataset. With these data, the z-scores of the cellular regulome enrichment scores were calculated and depicted in a heat map to compare relative regulatory activity between datasets. Regulon specificity scores were calculated using the ‘regulon_specificity_scores’ function from pySCENIC according to the ‘Cell_Class’ annotations.

### Identification of differentially expressed genes

Genes differentially expressed were calculated between N_SVZ vs T_SVZ, N_SVZ vs T_Mass, and T_SVZ vs T_Mass as a whole area. The R package LIBRA (v1.0.0) was used to perform zero-inflated regression analysis, as wrapper of the R library MAST^16^. Genes were defined as significantly differentially expressed at Benjamini–Hochberg correction FDR<0.05 and abs (log_2_(Fold Change))>0.3.

### Cell-cell interaction analysis

Intercellular communication network analysis was performed by using the standard workflow of the R package ‘CellChat’ (v1.4.0)^17^ with the CellChatDB.human database to assess the primary signaling inputs and outputs. Ligand-receptor analysis framework LIANA (v0.1.12)^18^, based on the consensus rank aggregate score calculated combining multiple algorithms as NATMI, iTalk, Connectome, SingleCellSignalR, and CellphoneDB, was used to detect interactions between microglia with other tumor cells in each area. MultiNicheNet (https://github.com/saeyslab/multinichenetr) package was used to find the differences in communication of microglia cells in T_SVZ and T_Mass region.

### Pathway and Macspectrum analyses

The functional annotation of the identified DEGs was performed using enrichGO from clusterProfiler R package^19^. Functional categories were selected based on FDR<0.05. MacSpectrum (V1.0.1)^20^ a tool that uses macrophage differentiation (MDI) and polarization indexes (MPI) previously generated using *in vitro* systems was utilized to further study activation gene signatures in microglia and MDM in all three areas. To functionally characterize MDM and microglia, we used the following parameters: “pre-activation” or “M0” cells (AMDI < 0, MPI < 0), “M1-transitional” (AMDI < 0, MPI > 0) or “M1-like” cells (AMDI > 0, MPI > 0), and “M2-like” cells (AMDI > 0, MPI < 0).

### Spatial transcriptomics

Sections of formalin-fixed paraffin-embeded (FFPE) tissues of 4 GBM (GBM4, GBM7, GBM8, and GBM9, with T_Mass and T_SVZ for each) were used per 10X Genomics *Visium Spatial Gene Expression for FFPE – Tissue Preparation Guide* (CG000408/Rev D) as follows: FFPE blocks were first faced and scored with a scalpel blade to isolate an area of tissue up to 6 x 6 mm, then chilled on an ice block for 20 minutes. Paraffin sections were cut at 5µm using a standard microtome, and floated on a 40℃ waterbath, containing purified water, to remove folds and wrinkles. Sections were carefully removed from the surface of the waterbath onto a Visium Spatial Gene Expression Slide within the fiducial frames, starting with the top frame. The microtome was cleaned with xylene substitute, alcohol, and RNAse Away, and a new blade was obtained between each block. Once all frames were filled, slides were placed in a slide rack in an oven at 42℃ for three hours, then stored overnight at room temperature within a slide box containing a desiccant packet.

After overnight drying, slides were deparaffinized, stained with Hematoxylin and Eosin (H&E), and imaged per 10X Genomics *Visium Spatial gene Expression for FFPE – Deparaffinization, H&E Staining, Imaging & Decrosslinking* (CG000409/Rev C) as follows:

i. for the slide deparaffinization, the following steps were taken: Xylene (3 changes, 5 minutes each), 100% ethanol (3 changes, 3 minutes each), 95% ethanol (2 changes, 3 minutes each), 85% ethanol (3 minutes), 70% ethanol (3 minutes), Purified water (1 minute).
ii. for the Hematoxylin and Eosin staining the following steps were taken: Mayers Hematoxylin (Millipore Sigma MHS16) – 3 minutes, Rinse in purified water (2 changes, 20 seconds and 10 seconds), Bluing Buffer (Fisher Scientific 6769001) – 1 minute, Rinse in purified water (5 dips plus 20 seconds), Alcoholic Eosin (Millipore Sigma HT110116) – 1 minute, Rinse in purified water (5 dips plus 20 seconds).

Slides were then coverslipped using 85% glycerol (Millipore Sigma 49781) and immediately imaged at 20X using the Leica Aperio AT2 slide digitizing system. Coverslips were removed and slides were carefully rinsed in purified water and allowed to air dry. Decrosslinking was performed with 0.1N HCl and TE Buffer (pH 9.0) to release RNA sequestered by formalin. Pairs of human transcriptome probes (PN-1000363, 10X Genomics) were hybridized to the RNA for 20 hours, and libraries were prepared following the manufacturer’s instructions (CG000407 Rev D). Libraries were sequenced on a Singular Genomics G4 following PCR to add G4-specific adapters. The PCR used 2 ng of library, 0.3 µM of each primer, and NEBNext Ultra II Q5 Master Mix (New England BioLabs) incubated at 98 °C for 2 min followed by 7 cycles of 98 °C for 20 sec, 57 °C for 30 sec, and 72 °C for 30 sec, with a final extension at 72 °C for 1 min. A similar protocol was used by 3D Genomics (https://3dgeno.com/) for the HNS1 sample (one of the two N_SVZ samples included in this study).

### 10X Genomics Visium analysis

Spatial transcriptomic data from the 4 GBM (GBM4, GBM7, GBM8, and GBM9) and from HNS1 were processed from FASTQ files and slides using 10x Genomics Space Ranger (v2.0.1). The GRCh38-2020-A reference transcriptome and the Human Transcriptome v1 Probe Set were used for alignment. Spatial data was then processed and visualized using Squidpy (v1.3.1)^21^ for Python-based analyses and Seurat (v4.9.9.9045)^5^ for R-based analyses. Downstream analysis was performed primarily in Python. Mitochondrial genes were filtered out, and SCTransform (v2)^22^ was used to correct the raw counts using Analytic Pearson Residuals. Mitochondrial genes were also filtered out prior to downstream analysis.

### Definition of malignant spots

Spatial copy number variation analysis was performed using the R packages SpatialInferCNV (v1.0.1)^23^ and InferCNV (v1.16.0) (https://github.com/broadinstitute/inferCNV) For each spot in each dataset, the raw gene expression counts were compiled and annotated by sample and tumor status. The N_SVZ data were used as a reference. The following parameters were used to generate the copy number variation heatmaps: cutoff=0.1, denoise=TRUE, HMM=FALSE.

### 10x Visium spot deconvolution

For cellular deconvolution, Cell2Location (v0.1.3)^24^ was used on all samples to determine the cell type abundances within each Visium spot. The number of cells within each spot was determined using Squidpy, and the average value was provided as a parameter to Cell2Location in order to perform deconvolution. Cell2Location was used to filter genes using the filter_genes function with the default parameters (cell_count_cutoff=5, cell_percentage_cutoff=0.03, nonz_mean_cutoff=1.12), resulting in quantities between 11,017 and 13,220 genes per dataset. We used Cell2Location’s RegressionModel to help map a posterior distribution of the estimated cell type abundance for each Cell Class in the single-cell data. This model was trained for 500 epochs. With the estimated cell-type abundances, a full Cell2Location model was trained for 30,000 epochs on the spatial data to estimate the deconvolved cell types in each Visium spot. For visualizations, Cell2Location’s plot_spatial function was used for each slide.

### 10X Genomics Visium sample clustering

Spatial data were clustered using Seurat (v4.9.9.9045)^5^ using the Leiden algorithm. Gene programs were defined per Leiden cluster by differential expression analysis based on the top 50 most differentially expressed genes by the Wilcoxon Rank Sum Test with a *p*<0.005.

### 10x Genomics Visium spot clustering and intracellular communication

The cell type proportions of each sample were computed using the cell type abundance figures from Cell2Location (v0.1.3)^24^. For each sample, the cell type abundance per spot was averaged and normalized, giving an overall cell type proportion for the entire sample. Non-negative matrix factorization was computed using Cell2Location to highlight cellular compartments. The run_colocation algorithm was run with default parameters, with the exception of n_fact, which was set to between 5 and 30 to explore a wide range of factors. The output data was reassembled into new plots to remove factors with no data. Cell2Location was used to identify cell-type specific expression of all genes in the datasets as a prerequisite for Node-Centric Expression Models. The Python library NCEM (v0.1.5)^25^ was used to infer cell-cell interaction and produce visualizations. In the network plots, edges are only plotted between sender and receiver cell types that share more than 150 differentially expressed genes. Edge thicknesses are proportional to the L1 norm of the vector of fold changes.

### Cancer stem-like cells establishment and propagation

Patient-derived cancer stem-like cells (CSCs) from the T_Mass and the T_SVZ tissues of the GBM patients included in this study were established as described previously^26^ and propagated *in vitro* using growth factor-enriched, serum-free cell culture medium based on Neurobasal™-A Medium, minus phenol red (Thermo Fisher Scientific, Cat. No. 12349015), N2 Supplement (Thermo Fisher Scientific, Cat. No. 17502048), B-27 Supplement, minus vitamin A (Thermo Fisher Scientific, Cat. No. 12587010), human bFGF (Thermo Fisher Scientific, Cat. No. PHG0261), human EGF (Thermo Fisher Scientific, Cat. No. PHG6045) and Pen/Strep/Glutamine (Thermo Fisher Scientific, Cat. No. 10378016).

### CD11b enrichment

Starting from briefly cultured cells obtained from the T_Mass and the T_SVZ of 4 GBM patients (GBM 4, 7, 17, and 23) included in this study, immunomagnetic microbeads decorated with recombinantly engineered antibody fragments for CD11b (Miltenyi Biotec, Cat. No. 130-049-601) were used with the MiniMACS Separation Unit (Miltenyi Biotec, Cat. No. 130-042-102), MACS MultiStand (Miltenyi Biotec, Cat. No. 130-042-303) and MS Columns (Miltenyi Biotec, Cat. No. 130-042-201) to enrich for the CD11b+ cell fraction. Specifically, cell numbers were counted and incubated with CD11b microbeads for 15 min at 4°C. Following centrifugation and resuspension, cells were run through the columns using the separation unit. Upon immunomagnetic separation, CD11b-enriched cells were plated at approximately 10^4^cells/cm^2^ in Macrophage Base Medium XF (PromoCell, Cat. No. C-28057) or M2-Macrophage Generation Medium XF (PromoCell, Cat. No. C-28056) prepared and supplemented with cytokines and mix as per the manufacturer’s instructions.

### IL-1β and Wnt-5a ELISA

Cell culture supernatants of 3×10^4^ cd11b-enriched tumor-associated macrophages (TAMs) isolated from the T_Mass and the T_SVZ of the 4 GBM patients (GBM4, 7, 17, and 23) and propagated *in vitro* using the M2-Macrophage Generation Medium XF (PromoCell, Cat. No. C-28056) were collected 48 hours after cell passaging or after 48 hours from treatment start and stored at −80C until the assay was performed. The Human IL-1 beta/IL-1F2 DuoSet ELISA 96-wells (R&D Systems, Cat. No. DY201-05) and the Wnt-5a ELISA kit (Human): 96 wells (Aviva Systems Biology, Cat. No. OKEH00723) were used as per the manufacturer’s recommendation. The experiments were performed in triplicates and repeated three times.

IL-1β and Wnt-5a ELISA were also performed on TAMs following treatment with Nidanilimab or Box5. Specifically, TAMs isolated from 2 GBM patients ((GBM4, and 7 with the T_Mass and the T_SVZ for each) were treated with the IL-1RAcP fully humanized monoclonal antibody Nidanilimab (Selleckchem, Cat. No. 300113) at 20 µg/ml or with the Wnt-5a antagonist Box5 (Selleckchem, Cat. No. P1216) at 100 µM for 48 hours. A total of 3×10^4^ TAMs were plated in triplicate per treatment condition and were treated for 48 hours with Nidanilimab or Box5 starting the day after plating. Control wells for each condition (patient and area) were also included in triplicate and the experiment was repeated three times. Cell culture supernatants were collected for ELISA.

### IL-1RAcP and Frizzled-3 immunofluorescence

CSCs were plated onto glass coverslips coated with Matrigel (Fisher Scientific, Cat. No. CB-40234A) overnight. Cells were washed and fixed in 4% Formaldehyde (Millipore Sigma, Cat. No. M2-01-04) for 10 minutes at room temperature following permeabilization with 0.02% Triton X-100 (Millipore Sigma, Cat. No. T8787). After 1 hour of blocking with 5% goat serum (Millipore Sigma, Cat. No. G9023), cells were processed for immunofluorescence using an IL-1RAcP monoclonal antibody (Abnova, Cat. No. H00003556-M03) and the FZD3 polyclonal antibody (Millipore Sigma, Cat. No. SAB4503171) at the recommended dilution of 1:200 at 4°C overnight. The next day, cells were washed with 1X phosphate buffer saline (PBS) (Millipore Sigma, Cat. No. P2272) and incubated for 1 hour at room temperature with Alexa Fluor™488 (Thermo Fisher Scientific, Cat. No. A32731 and A32723) used at 1:500. Cells were then counterstained with DAPI (Thermo Fisher Scientific, Cat. No. 62248) used at 0.5 µg/ml for 10 minutes. After washing with PBS, coverslips were mounted onto microscope slides in mounting media. Slides were imaged in a ZEISS LSM 900 Confocal Microscope. Image exporting to TIFF files was obtained using ZEN (blue edition). The same steps were performed for IL-1RAcP Immunostaining of TAMs.

### *In vitro* treatment assay

To evaluate the efficacy of Nidanilimab, a IL-1RAcP fully humanized monoclonal antibody (Selleckchem, Cat. No. 300113), CSCs isolated from 3 GBM patients (T_Mass and T_SVZ for each) and propagated in serum-free conditions as described above were treated with IL-1β and Nidanilimab. Briefly, IL-1β treatment was performed by adding the recombinant human IL-1β protein (R&D Systems, Cat. No. 201-LB/CF) to growth factor-enriched, serum-free cell culture medium at a final concentration of 100ng/ml. Nidanilimab was used at 20 µg/ml. A total of 2.5×10^5^ CSC were plated in triplicate per treatment condition and were treated for 48 hours with IL-1β alone or with IL-1β and Nidanilimab starting the day after plating. The *in vitro* proliferative potential of CSCs was evaluated as previously described^27^. Non-linear regression analysis was performed to assess the proliferative potential.

To evaluate the efficacy of Box5, a Wnt-5a antagonist (Selleckchem, Cat. No. P1216), CSCs isolated from 3 GBM patients (T_Mass and T_SVZ for each) and propagated in serum-free conditions as described above were exposed to conditioned medium of matched TAMs treated *in vitro* with Box5 for 48 hours. Briefly, a total of 3×10^4^ matched TAMs were plated in triplicate per treatment condition. Control wells for each condition (patient and area) were also included in triplicate. One day after plating, cells were treated with Box5 at 100 µM for 48 hours. 1×10^4^ cells/cm^2^ CSCs plated in triplicates on transwell polycarbonate membranes with cell culture inserts (Millipore Sigma, Cat. No. CLS3428) pre-coated with Matrigel (Corning, Cat. No. 356234) diluted 1:50 in cell culture medium were exposed for 10-14 days to the conditioned medium of matched TAMs, treated and control. To quantify cell invasion, cells on the inside of the transwell membranes were removed using cotton swabs, and those invading the lower surface of the membranes were stained with 0.25% crystal violet (Millipore Sigma, Cat. No. 61135) for 10 minutes. After washing and drying, membranes were removed from the inserts and imaged by microscopy. Crystal violet was then eluted from the membranes by adding 500 µl of 33% v/v acetic acid (Millipore Sigma, Cat. No. AX0073-75) solution and shaking for 10 minutes. The eluted crystal violet was then transferred to a 96-multiwell plate (100 µl/well). Absorbance at 590 nm was measured using a plate reader. The experiment was repeated three times.

### Survival analysis

Kaplan Meier survival curves were obtained by plotting the *IL1RAP* and *FZD3* expression levels and patient survival using data of the The Cancer Genome Atlas and Rembrandt datasets. The GlioVis data portal^28^ was used to correlate and determine the statistical significance of the *IL1RAP* and *FZD3* expression levels with patient survival.

## References

1 Stupp, R. et al. Radiotherapy plus concomitant and adjuvant temozolomide for glioblastoma. N Engl J Med 352, 987–996, doi:10.1056/NEJMoa043330 (2005).

2 Wen, P. Y. & Kesari, S. Malignant gliomas in adults. N Engl J Med 359, 492–507, doi:10.1056/NEJMra0708126 (2008).

3 Ostrom, Q. T. et al. CBTRUS Statistical Report: Primary Brain and Other Central Nervous System Tumors Diagnosed in the United States in 2016-2020. Neuro Oncol 25, iv1-iv99, doi:10.1093/neuonc/noad149 (2023).

4 Piccirillo, S. G. et al. Contributions to drug resistance in glioblastoma derived from malignant cells in the sub-ependymal zone. Cancer Res 75, 194–202, doi:10.1158/0008-5472.CAN-13-3131 (2015).

5 Spiteri, I. et al. Evolutionary dynamics of residual disease in human glioblastoma. Ann Oncol 30, 456–463, doi:10.1093/annonc/mdy506 (2019).

6 Markovic, D. S. et al. Gliomas induce and exploit microglial MT1-MMP expression for tumor expansion. Proc Natl Acad Sci U S A 106, 12530–12535, doi:10.1073/pnas.0804273106 (2009).

7 Coniglio, S. J. et al. Microglial stimulation of glioblastoma invasion involves epidermal growth factor receptor (EGFR) and colony stimulating factor 1 receptor (CSF-1R) signaling. Mol Med 18, 519–527, doi:10.2119/molmed.2011.00217 (2012).

8 Pyonteck, S. M. et al. CSF-1R inhibition alters macrophage polarization and blocks glioma progression. Nat Med 19, 1264–1272, doi:10.1038/nm.3337 (2013).

9 Zhou, W. et al. Periostin secreted by glioblastoma stem cells recruits M2 tumour-associated macrophages and promotes malignant growth. Nat Cell Biol 17, 170–182, doi:10.1038/ncb3090 (2015).

10 Gabrusiewicz, K. et al. Glioblastoma-infiltrated innate immune cells resemble M0 macrophage phenotype. JCI Insight 1, doi:10.1172/jci.insight.85841 (2016).

11 Bowman, R. L. et al. Macrophage Ontogeny Underlies Differences in Tumor-Specific Education in Brain Malignancies. Cell Rep 17, 2445–2459, doi:10.1016/j.celrep.2016.10.052 (2016).

12 Shi, Y. et al. Tumour-associated macrophages secrete pleiotrophin to promote PTPRZ1 signalling in glioblastoma stem cells for tumour growth. Nat Commun 8, 15080, doi:10.1038/ncomms15080 (2017).

13 Wang, Q. et al. Tumor Evolution of Glioma-Intrinsic Gene Expression Subtypes Associates with Immunological Changes in the Microenvironment. Cancer Cell 32, 42–56 e46, doi:10.1016/j.ccell.2017.06.003 (2017).

14 Darmanis, S. et al. Single-Cell RNA-Seq Analysis of Infiltrating Neoplastic Cells at the Migrating Front of Human Glioblastoma. Cell Rep 21, 1399–1410, doi:10.1016/j.celrep.2017.10.030 (2017).

15 Muller, S. et al. Single-cell profiling of human gliomas reveals macrophage ontogeny as a basis for regional differences in macrophage activation in the tumor microenvironment. Genome Biol 18, 234, doi:10.1186/s13059-017-1362-4 (2017).

16 Neftel, C. et al. An Integrative Model of Cellular States, Plasticity, and Genetics for Glioblastoma. Cell 178, 835–849 e821, doi:10.1016/j.cell.2019.06.024 (2019).

17 Takenaka, M. C. et al. Control of tumor-associated macrophages and T cells in glioblastoma via AHR and CD39. Nat Neurosci 22, 729–740, doi:10.1038/s41593-019-0370-y (2019).

18 Kaffes, I. et al. Human Mesenchymal glioblastomas are characterized by an increased immune cell presence compared to Proneural and Classical tumors. Oncoimmunology 8, e1655360, doi:10.1080/2162402X.2019.1655360 (2019).

19 Chen, P. et al. Circadian Regulator CLOCK Recruits Immune-Suppressive Microglia into the GBM Tumor Microenvironment. Cancer Discov 10, 371–381, doi:10.1158/2159-8290.CD-19-0400 (2020).

20 Friebel, E. et al. Single-Cell Mapping of Human Brain Cancer Reveals Tumor-Specific Instruction of Tissue-Invading Leukocytes. Cell 181, 1626–1642 e1620, doi:10.1016/j.cell.2020.04.055 (2020).

21 Klemm, F. et al. Interrogation of the Microenvironmental Landscape in Brain Tumors Reveals Disease-Specific Alterations of Immune Cells. Cell 181, 1643–1660 e1617, doi:10.1016/j.cell.2020.05.007 (2020).

22 Landry, A. P., Balas, M., Alli, S., Spears, J. & Zador, Z. Distinct regional ontogeny and activation of tumor associated macrophages in human glioblastoma. Sci Rep 10, 19542, doi:10.1038/s41598-020-76657-3 (2020).

23 Ochocka, N. et al. Single-cell RNA sequencing reveals functional heterogeneity of glioma-associated brain macrophages. Nat Commun 12, 1151, doi:10.1038/s41467-021-21407-w (2021).

24 Pombo Antunes, A. R., et al. Single-cell profiling of myeloid cells in glioblastoma across species and disease stage reveals macrophage competition and specialization. Nat Neurosci 24, 595–610, doi:10.1038/s41593-020-00789-y (2021).

25 Ravi, V. M. et al. T-cell dysfunction in the glioblastoma microenvironment is mediated by myeloid cells releasing interleukin-10. Nat Commun 13, 925, doi:10.1038/s41467-022-28523-1 (2022).

26 Ravi, V. M. et al. Spatially resolved multi-omics deciphers bidirectional tumor-host interdependence in glioblastoma. Cancer Cell 40, 639–655 e613, doi:10.1016/j.ccell.2022.05.009 (2022).

27 Yin, W. et al. A map of the spatial distribution and tumour-associated macrophage states in glioblastoma and grade 4 IDH-mutant astrocytoma. J Pathol 258, 121–135, doi:10.1002/path.5984 (2022).

28 Kim, H. J. et al. Blood monocyte-derived CD169(+) macrophages contribute to antitumor immunity against glioblastoma. Nat Commun 13, 6211, doi:10.1038/s41467-022-34001-5 (2022).

29 Garcia-Montano, L. A. et al. Dissecting Intra-tumor Heterogeneity in the Glioblastoma Microenvironment Using Fluorescence-Guided Multiple Sampling. Mol Cancer Res, OF1-OF13, doi:10.1158/1541-7786.MCR-23-0048 (2023).

30 Brennan, C. W. et al. The somatic genomic landscape of glioblastoma. Cell 155, 462–477, doi:10.1016/j.cell.2013.09.034 (2013).

31 Ceccarelli, M. et al. Molecular Profiling Reveals Biologically Discrete Subsets and Pathways of Progression in Diffuse Glioma. Cell 164, 550–563, doi:10.1016/j.cell.2015.12.028 (2016).

32 Durante, M. A. et al. Single-cell analysis reveals new evolutionary complexity in uveal melanoma. Nat Commun 11, 496, doi:10.1038/s41467-019-14256-1 (2020).

33 Couturier, C. P. et al. Glioblastoma scRNA-seq shows treatment-induced, immune-dependent increase in mesenchymal cancer cells and structural variants in distal neural stem cells. Neuro Oncol 24, 1494–1508, doi:10.1093/neuonc/noac085 (2022).

34 Lange, M. et al. CellRank for directed single-cell fate mapping. Nat Methods 19, 159–170, doi:10.1038/s41592-021-01346-6 (2022).

35 Weiler, P., Lange, M., Klein, M., Pe’er, D. & Theis, F. J. Unified fate mapping in multiview single-cell data. bioRxiv, 10.1101/2023.07.19.549685 (2023).

36 Aibar, S. et al. SCENIC: single-cell regulatory network inference and clustering. Nat Methods 14, 1083–1086, doi:10.1038/nmeth.4463 (2017).

37 Van de Sande, B. et al. A scalable SCENIC workflow for single-cell gene regulatory network analysis. Nat Protoc 15, 2247–2276, doi:10.1038/s41596-020-0336-2 (2020).

38 Tome-Garcia, J. et al. Analysis of chromatin accessibility uncovers TEAD1 as a regulator of migration in human glioblastoma. Nat Commun 9, 4020, doi:10.1038/s41467-018-06258-2 (2018).

39 Barrette, A. M. et al. Anti-invasive efficacy and survival benefit of the YAP-TEAD inhibitor verteporfin in preclinical glioblastoma models. Neuro Oncol 24, 694–707, doi:10.1093/neuonc/noab244 (2022).

40 Siebzehnrubl, F. A. et al. The ZEB1 pathway links glioblastoma initiation, invasion and chemoresistance. EMBO Mol Med 5, 1196–1212, doi:10.1002/emmm.201302827 (2013).

41 Singh, D. K. et al. Oncogenes Activate an Autonomous Transcriptional Regulatory Circuit That Drives Glioblastoma. Cell Rep 18, 961–976, doi:10.1016/j.celrep.2016.12.064 (2017).

42 Chandra, A. et al. Clonal ZEB1-Driven Mesenchymal Transition Promotes Targetable Oncologic Antiangiogenic Therapy Resistance. Cancer Res 80, 1498–1511, doi:10.1158/0008-5472.CAN-19-1305 (2020).

43 Mossi Albiach, A. J. J.; Kapustova, I.; Kvedaraite, E.; Codeluppi, S.; Munting, J. B.; Borm, L. E.; Jacobsen, J. K.; Shamikh, A.; Persson, O.; Linnarsson, S. Glioblastoma is spatially organized by neurodevelopmental programs and a glial-like wound healing response. bioRxiv, 10.1101/2023.09.01.555882 (2023).

44 Jing, S. et al. Expression of TCF7L2 in Glioma and Its Relationship With Clinicopathological Characteristics and Patient Overall Survival. Front Neurol 12, 627431, doi:10.3389/fneur.2021.627431 (2021).

45 Nixon, B. G. et al. Tumor-associated macrophages expressing the transcription factor IRF8 promote T cell exhaustion in cancer. Immunity 55, 2044–2058 e2045, doi:10.1016/j.immuni.2022.10.002 (2022).

46 Masuda, T. et al. IRF8 is a critical transcription factor for transforming microglia into a reactive phenotype. Cell Rep 1, 334–340, doi:10.1016/j.celrep.2012.02.014 (2012).

47 Li, C. et al. Single cell transcriptomics based-MacSpectrum reveals novel macrophage activation signatures in diseases. JCI Insight 5, doi:10.1172/jci.insight.126453 (2019).

48 Suva, M. L. et al. Reconstructing and reprogramming the tumor-propagating potential of glioblastoma stem-like cells. Cell 157, 580–594, doi:10.1016/j.cell.2014.02.030 (2014).

49 Stevanovic, M. et al. SOX Transcription Factors as Important Regulators of Neuronal and Glial Differentiation During Nervous System Development and Adult Neurogenesis. Front Mol Neurosci 14, 654031, doi:10.3389/fnmol.2021.654031 (2021).

50 Comba, A. et al. Spatiotemporal analysis of glioma heterogeneity reveals COL1A1 as an actionable target to disrupt tumor progression. Nat Commun 13, 3606, doi:10.1038/s41467-022-31340-1 (2022).

51 Sarkar, S. & Yong, V. W. Inflammatory cytokine modulation of matrix metalloproteinase expression and invasiveness of glioma cells in a 3-dimensional collagen matrix. J Neurooncol 91, 157–164, doi:10.1007/s11060-008-9695-1 (2009).

52 Li, Y., Wang, L., Pappan, L., Galliher-Beckley, A. & Shi, J. IL-1beta promotes stemness and invasiveness of colon cancer cells through Zeb1 activation. Mol Cancer 11, 87, doi:10.1186/1476-4598-11-87 (2012).

53 Carmi, Y. et al. The role of IL-1beta in the early tumor cell-induced angiogenic response. J Immunol 190, 3500–3509, doi:10.4049/jimmunol.1202769 (2013).

54 Herting, C. J. et al. Tumour-associated macrophage-derived interleukin-1 mediates glioblastoma-associated cerebral oedema. Brain 142, 3834–3851, doi:10.1093/brain/awz331 (2019).

55 Bayik, D. et al. Myeloid-Derived Suppressor Cell Subsets Drive Glioblastoma Growth in a Sex-Specific Manner. Cancer Discov 10, 1210–1225, doi:10.1158/2159-8290.CD-19-1355 (2020).

56 Liu, H. et al. Pro-inflammatory and proliferative microglia drive progression of glioblastoma. Cell Rep 36, 109718, doi:10.1016/j.celrep.2021.109718 (2021).

57 Kai, K. et al. Macrophage/microglia-derived IL-1beta induces glioblastoma growth via the STAT3/NF-kappaB pathway. Hum Cell 35, 226–237, doi:10.1007/s13577-021-00619-8 (2022).

58 Chen, Z. et al. A paracrine circuit of IL-1beta/IL-1R1 between myeloid and tumor cells drives genotype-dependent glioblastoma progression. J Clin Invest, doi:10.1172/JCI163802 (2023).

59 Kamino, M. et al. Wnt-5a signaling is correlated with infiltrative activity in human glioma by inducing cellular migration and MMP-2. Cancer Sci 102, 540–548, doi:10.1111/j.1349-7006.2010.01815.x (2011).

60 Hu, B. et al. Epigenetic Activation of WNT5A Drives Glioblastoma Stem Cell Differentiation and Invasive Growth. Cell 167, 1281–1295 e1218, doi:10.1016/j.cell.2016.10.039 (2016).

61 Binda, E. et al. Wnt5a Drives an Invasive Phenotype in Human Glioblastoma Stem-like Cells. Cancer Res 77, 996–1007, doi:10.1158/0008-5472.CAN-16-1693 (2017).

62 Liu, G. et al. Daam1 activates RhoA to regulate Wnt5a-induced glioblastoma cell invasion. Oncol Rep 39, 465–472, doi:10.3892/or.2017.6124 (2018).

63 Trivieri, N. et al. Growth factor independence underpins a paroxysmal, aggressive Wnt5a(High)/EphA2(Low) phenotype in glioblastoma stem cells, conducive to experimental combinatorial therapy. J Exp Clin Cancer Res 41, 139, doi:10.1186/s13046-022-02333-1 (2022).

64 Dijksterhuis, J. P. et al. High levels of WNT-5A in human glioma correlate with increased presence of tumor-associated microglia/monocytes. Exp Cell Res 339, 280–288, doi:10.1016/j.yexcr.2015.10.022 (2015).

65 Fischer, D. S., Schaar, A. C. & Theis, F. J. Modeling intercellular communication in tissues using spatial graphs of cells. Nat Biotechnol 41, 332–336, doi:10.1038/s41587-022-01467-z (2023).

66 Pietras, A. et al. Osteopontin-CD44 signaling in the glioma perivascular niche enhances cancer stem cell phenotypes and promotes aggressive tumor growth. Cell Stem Cell 14, 357–369, doi:10.1016/j.stem.2014.01.005 (2014).

67 Szulzewsky, F. et al. Glioma-associated microglia/macrophages display an expression profile different from M1 and M2 polarization and highly express Gpnmb and Spp1. PLoS One 10, e0116644, doi:10.1371/journal.pone.0116644 (2015).

68 Chen, P. et al. Symbiotic Macrophage-Glioma Cell Interactions Reveal Synthetic Lethality in PTEN-Null Glioma. Cancer Cell 35, 868–884 e866, doi:10.1016/j.ccell.2019.05.003 (2019).

69 Abdelfattah, N. et al. Single-cell analysis of human glioma and immune cells identifies S100A4 as an immunotherapy target. Nat Commun 13, 767, doi:10.1038/s41467-022-28372-y (2022).

70 He, C. et al. Single-Cell Transcriptomic Analysis Revealed a Critical Role of SPP1/CD44-Mediated Crosstalk Between Macrophages and Cancer Cells in Glioma. Front Cell Dev Biol 9, 779319, doi:10.3389/fcell.2021.779319 (2021).

71 Robbrecht, D. et al. First-in-human phase 1 dose-escalation study of CAN04, a first-in-class interleukin-1 receptor accessory protein (IL1RAP) antibody in patients with solid tumours. Br J Cancer 126, 1010–1017, doi:10.1038/s41416-021-01657-7 (2022).

72 Paulus, A. C. S.; Zvirbule, Z.; Paz-Ares, L.; Awada, A.; Garcia-Ribas, I.; Losic, N.; Zemaitis, M. Phase 1/2a trial of nadunolimab, a first-in-class fully humanized monoclonal antibody against IL1RAP, in combination with cisplatin and gemcitabine (CG) in patients with non-small cell lung cancer (NSCLC). J Clin Oncol. 40: 9020, doi:10.1200/JCO.2022.40.16_suppl.9020 (2022).

73 Van Cutsem, E. L. E. R.; Ochsenreither, S.; Zvirbule, Z.; Ivanauskas, A.; Arnold, D.; Baltruskeviciene, E.; Pfeiffer, P.; Yachnin, J.; Garcia-Carbonero, R.; Greil, R.; Jungels, C.; Poulsen, L.; Awada, A.; Garcia-Ribas, I.; Losic, N.; Collignon, J. Phase 1/2a trial of nadunolimab, a first-in-class fully humanized monoclonal antibody against IL1RAP, in combination with gemcitabine and nab-paclitaxel (GN) in patients with pancreatic adenocarcinoma (PDAC). J Clin Oncol. 40: 4141, doi:10.1200/JCO.2022.40.16_suppl.4141 (2022).

74 Miller, T. E. et al. Programs, Origins, and Niches of Immunomodulatory Myeloid Cells in Gliomas. bioRxiv, doi:10.1101/2023.10.24.563466 (2023).

75 Ruiz-Moreno, C. et al. Harmonized single-cell landscape, intercellular crosstalk and tumor architecture of glioblastoma bioRxiv, doi:10.1101/2022.08.27.505439 (2022).

76 Bartkowiak, T. et al. An immunosuppressed microenvironment distinguishes lateral ventricle-contacting glioblastomas. JCI Insight 8, doi:10.1172/jci.insight.160652 (2023).

77 Masuda, T., Sankowski, R., Staszewski, O. & Prinz, M. Microglia Heterogeneity in the Single-Cell Era. Cell Rep 30, 1271–1281, doi:10.1016/j.celrep.2020.01.010 (2020).

78 Keane, L., Cheray, M., Blomgren, K. & Joseph, B. Multifaceted microglia - key players in primary brain tumour heterogeneity. Nat Rev Neurol 17, 243–259, doi:10.1038/s41582-021-00463-2 (2021).

79 Sankowski, R. et al. Mapping microglia states in the human brain through the integration of high-dimensional techniques. Nat Neurosci 22, 2098–2110, doi:10.1038/s41593-019-0532-y (2019).

80 Bowman, R. L., Wang, Q., Carro, A., Verhaak, R. G. & Squatrito, M. GlioVis data portal for visualization and analysis of brain tumor expression datasets. Neuro Oncol 19, 139-141, doi:10.1093/neuonc/now247 (2017).

## References

1 Cingolani, P. et al. A program for annotating and predicting the effects of single nucleotide polymorphisms, SnpEff: SNPs in the genome of Drosophila melanogaster strain w1118; iso-2; iso-3. Fly (Austin) 6, 80-92, doi:10.4161/fly.19695 (2012).

2 Durante, M. A. et al. Single-cell analysis reveals new evolutionary complexity in uveal melanoma. Nat Commun 11, 496, doi:10.1038/s41467-019-14256-1 (2020).

3 Zheng, G. X. et al. Massively parallel digital transcriptional profiling of single cells. Nat Commun 8, 14049, doi:10.1038/ncomms14049 (2017).

4 Frankish, A. et al. Gencode 2021. Nucleic Acids Res 49, D916–D923, doi:10.1093/nar/gkaa1087 (2021).

5 Hao, Y. et al. Integrated analysis of multimodal single-cell data. Cell 184, 3573–3587 e3529, doi:10.1016/j.cell.2021.04.048 (2021).

6 Germain, P. L., Lun, A., Garcia Meixide, C., Macnair, W. & Robinson, M. D. Doublet identification in single-cell sequencing data using scDblFinder. F1000Res 10, 979, doi:10.12688/f1000research.73600.2 (2021).

7 Hafemeister, C. & Satija, R. Normalization and variance stabilization of single-cell RNA-seq data using regularized negative binomial regression. Genome Biol 20, 296, doi:10.1186/s13059-019-1874-1 (2019).

8 Korsunsky, I. et al. Fast, sensitive and accurate integration of single-cell data with Harmony. Nat Methods 16, 1289–1296, doi:10.1038/s41592-019-0619-0 (2019).

9 Ianevski, A., Giri, A. K. & Aittokallio, T. Fully-automated and ultra-fast cell-type identification using specific marker combinations from single-cell transcriptomic data. Nat Commun 13, 1246, doi:10.1038/s41467-022-28803-w (2022).

10 Neftel, C. et al. An Integrative Model of Cellular States, Plasticity, and Genetics for Glioblastoma. Cell 178, 835–849 e821, doi:10.1016/j.cell.2019.06.024 (2019).

11 Weiler, P., Lange, M., Klein, M., Pe’er, D. & Theis, F. J. Unified fate mapping in multiview single-cell data. bioRxiv, 10.1101/2023.07.19.549685 (2023).

12 Lange, M. et al. CellRank for directed single-cell fate mapping. Nat Methods 19, 159–170, doi:10.1038/s41592-021-01346-6 (2022).

13 van Dijk, D. et al. Recovering Gene Interactions from Single-Cell Data Using Data Diffusion. Cell 174, 716–729 e727, doi:10.1016/j.cell.2018.05.061 (2018).

14 Reuter, B., Fackeldey, K. & Weber, M. Generalized Markov modeling of nonreversible molecular kinetics. J Chem Phys 150, 174103, doi:10.1063/1.5064530 (2019).

15 Van de Sande, B. et al. A scalable SCENIC workflow for single-cell gene regulatory network analysis. Nat Protoc 15, 2247–2276, doi:10.1038/s41596-020-0336-2 (2020).

16 Squair, J. W. et al. Confronting false discoveries in single-cell differential expression. Nat Commun 12, 5692, doi:10.1038/s41467-021-25960-2 (2021).

17 Jin, S. et al. Inference and analysis of cell-cell communication using CellChat. Nat Commun 12, 1088, doi:10.1038/s41467-021-21246-9 (2021).

18 Dimitrov, D. et al. Comparison of methods and resources for cell-cell communication inference from single-cell RNA-Seq data. Nat Commun 13, 3224, doi:10.1038/s41467-022-30755-0 (2022).

19 Yu, G., Wang, L. G., Han, Y. & He, Q. Y. clusterProfiler: an R package for comparing biological themes among gene clusters. OMICS 16, 284–287, doi:10.1089/omi.2011.0118 (2012).

20 Li, C. et al. Single cell transcriptomics based-MacSpectrum reveals novel macrophage activation signatures in diseases. JCI Insight 5, doi:10.1172/jci.insight.126453 (2019).

21 Palla, G. et al. Squidpy: a scalable framework for spatial omics analysis. Nat Methods 19, 171–178, doi:10.1038/s41592-021-01358-2 (2022).

22 Choudhary, S. & Satija, R. Comparison and evaluation of statistical error models for scRNA-seq. Genome Biol 23, 27, doi:10.1186/s13059-021-02584-9 (2022).

23 Erickson, A. et al. Spatially resolved clonal copy number alterations in benign and malignant tissue. Nature 608, 360–367, doi:10.1038/s41586-022-05023-2 (2022).

24 Kleshchevnikov, V. et al. Cell2location maps fine-grained cell types in spatial transcriptomics. Nat Biotechnol 40, 661–671, doi:10.1038/s41587-021-01139-4 (2022).

25 Fischer, D. S., Schaar, A. C. & Theis, F. J. Modeling intercellular communication in tissues using spatial graphs of cells. Nat Biotechnol 41, 332–336, doi:10.1038/s41587-022-01467-z (2023).

26 Piccirillo, S. G. et al. Contributions to drug resistance in glioblastoma derived from malignant cells in the sub-ependymal zone. Cancer Res 75, 194–202, doi:10.1158/0008-5472.CAN-13-3131 (2015).

27 Piccirillo, S. G. et al. Bone morphogenetic proteins inhibit the tumorigenic potential of human brain tumour-initiating cells. Nature 444, 761–765, doi:10.1038/nature05349 (2006).

28 Bowman, R. L., Wang, Q., Carro, A., Verhaak, R. G. & Squatrito, M. GlioVis data portal for visualization and analysis of brain tumor expression datasets. Neuro Oncol 19, 139-141, doi:10.1093/neuonc/now247 (2017).

