## Supplementary material for "Single-nucleus and spatial landscape of the sub-ventricular zone in human glioblastoma": Suppl. Information

### Supplementary Information

**Supplementary Figure 1.** Violin plots of the number of genes detected and UMI in each GBM patient and each area............................................................................................................ 4-5
**Supplementary Figure 2.** Copy-number variations for all GBM patients by area.................... 5-7
**Supplementary Table 4.** List of differentially expressed genes between N_SVZ *versus* T_Mass, N_SVZ *versus* T_SVZ and T_Mass *versus* T_SVZ……………................................... Appendix 1
**Supplementary Table 5.** List of differentially expressed genes in microglia between N_SVZ *versus* T_Mass, N_SVZ *versus* T_SVZ and T_Mass *versus* T_SVZ…………….......... Appendix 2
**Supplementary Figure 8.** Gene expression signature of 'homeostatic', 'activated', and 'inflammatory' microglia in the N_SVZ, T_Mass, and T_SVZ………………………………………11
**Supplementary Figure 9.** Tumor cell purity in the T_Mass and the T_SVZ of the 4 GBM patients…………………………………………………………………………………………. 11
**Supplementary Figure 10.** Cell type composition in the T_Mass and the T_SVZ of the 4 GBM patients…………………………………………………………………………………….…11-12
**Supplementary Figure 11.** Spatial frequency correlation between microglia and the other cell types in the T_Mass and the T_SVZ…………………………………………………...…….12-14
**Supplementary Figure 12.** Patterns of spatial dependencies among cell types in the T_Mass and T_SVZ of the 4 GBM patients and in the HNS1 sample…………………………………..14-16
**Supplementary Figure 17.** IL-1RAcP immunofluorescence of TAMs isolated from the T_Mass and the T_SVZ………………………..…………………………………………….…..…………..18
**Supplementary Figure 18.** Transwell assays of CSCs isolated from the T_Mass and the T_SVZ of GBM17 and exposed to conditioned medium of matched TAMs treated with Box5…………….18
**References**……………………………………………………………………………………...19

#### Supplementary Table 1. Clinical and molecular information of GBM patients.

| Patient ID | WHO Diagnosis | Tumor type | Sex | Age at Initial Diagnosis | Tumor Location | Prior surgery | MGMT | IDH 1/2 | Morphology |
|---|---|---|---|---|---|---|---|---|---|
| 4 | Glioblastoma | Primary | F | 83 | Left frontal | N | methylated | WT | |
| 7 | Glioblastoma | Primary | M | 77 | Right temporal | N | methylated | WT | |
| 8 | Glioblastoma | Primary | M | 64 | Right frontal | N | methylated | WT | giant cell |
| 9 | Breast cancer and Glioblastoma | Primary | F | 71 | Right parietal | N | unmethylated | WT | |
| 10 | Glioblastoma | Primary | M | 64 | Left temporal | N | methylated | WT | |
| 12 | Glioblastoma | Primary | F | 66 | Right parietotemporal | N | unmethylated | WT | |
| 14 | Glioblastoma | Primary | M | 61 | Right parietotemporal | N | low methylation | WT | |
| 16 | Glioblastoma | Primary | M | 77 | Right parietotemporal | N | methylated | WT | |
| 17 | Glioblastoma | Primary | F | 56 | Left frontal | N | unmethylated (?) | WT | |
| 18 | Glioblastoma | Primary | F | 78 | Right parietal lesion with subependymal spread and a satellite lesion in right temporal insula | N | NA | WT | multifocal |
| 20 | Progression from low-grade glioma | Secondary | M | 71 | Left deep posterior temporal | N | unmethylated | WT | |
| 22 | Glioblastoma | Primary | F | 80 | Right temporal | N | methylated | WT | |
| MIS1 | Glioblastoma | Recurrence | M | 21 | Left temporal, insular | Y | unmethylated | WT | |
| MIS2 | Glioblastoma | Primary | F | 73 | Right frontotemporal, insular | N | unmethylarted | WT | |
| MIS3 | Glioblastoma | Primary | F | 51 | Left temporal, insular | N | methylated | mut | |

The table summarizes the patient clinical and molecular characteristics. Abbreviations: MGMT, O(6)-methylguanine-DNA methyl transferase; NA, not available; IDH 1/2, Isocitrate Dehydrogenase (NADP(+)) 1/2; WT, wild-type.

**Supplementary Table 2. Molecular classification of GBM patients.** The table summarizes the status of the genetic drivers of GBM, as described by Verhaak *et al.*[1]. Only the results of the T_Mass samples of the analyzed patients are shown here. Abbreviations: cn, copy number; ge, gene expression; mut, mutation.

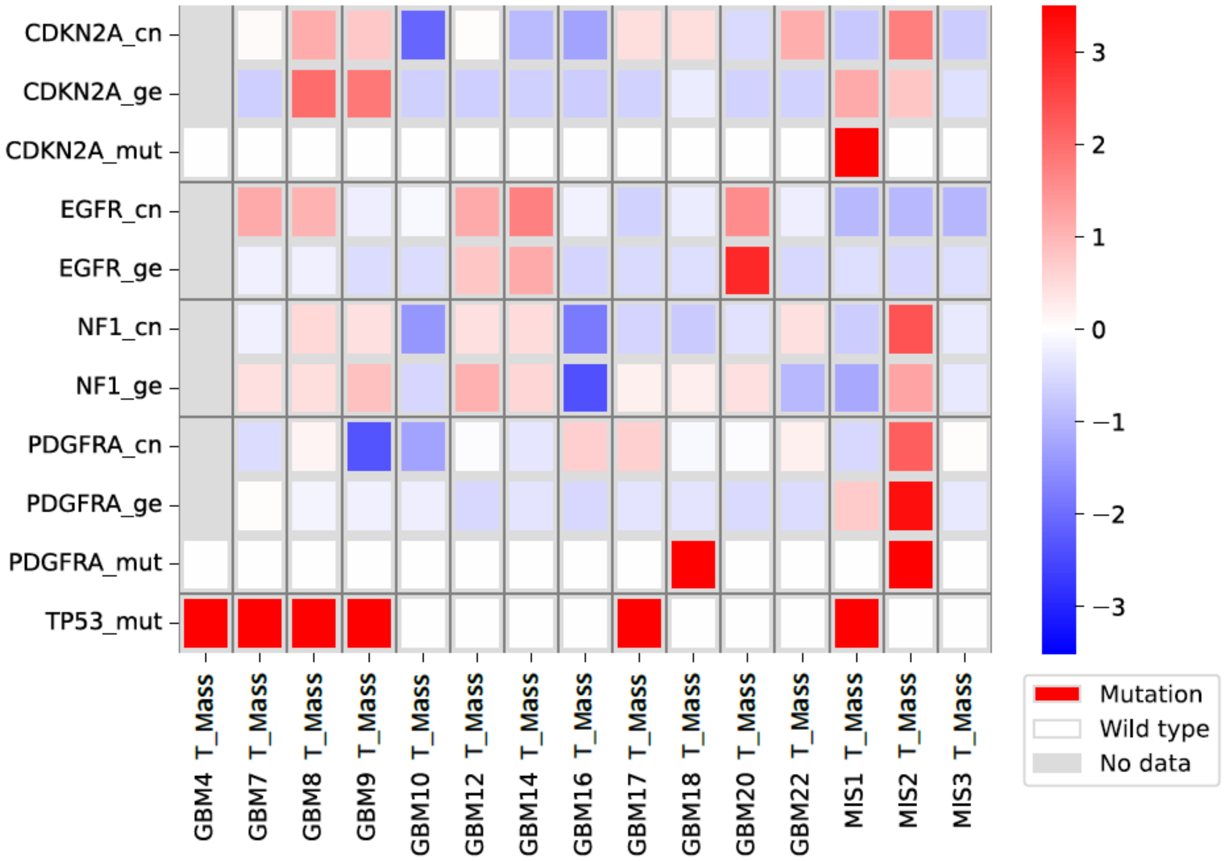

**Supplementary Table 3. Number of cells/nuclei obtained from each area of the 15 GBM patients and 2 controls.** The table shows the number of cells/nuclei obtained from each area. Briefly, Tumor mass and SVZ samples were collected from 15 GBM patients (GBM 4, 7, 8, 9, 10, 12, 14, 16, 17, 18, 20, 22, MIS1, MIS2, MIS3). Histologically normal SVZ samples were collected from two individuals: one SVZ was collected as postmortem tissue (HNS1) and the other during tumor surgical resection (HNS2). GBM4 samples were the only ones for which cells were obtained for single-cell RNA sequencing.

| GBM ID | Area | Number of tot. cells/nuclei per sample ($x10^5$) |
|---|---|---|
| 4 | Tumor mass | 20.2 |
| | SVZ | 18.1 |
| 7 | Tumor mass | 0.54 |
| | SVZ | 0.44 |
| 8 | Tumor mass | 1.07 |
| | SVZ | 1.4 |
| 9 | Tumor mass | 0.26 |
| | SVZ | 1.8 |
| 10 | Tumor mass | 2.1 |
| | SVZ | 2.27 |
| 12 | Tumor mass | 1.98 |
| | SVZ | 3.5 |
| 14 | Tumor mass | 0.782 |
| | SVZ | 10 |
| 16 | Tumor mass | 15 |
| | SVZ | 44 |
| 17 | Tumor mass | 5.28 |
| | SVZ | 1.89 |
| 18 | Tumor mass | 5.43 |
| | SVZ | 1.35 |
| 20 | Tumor mass | 13.6 |
| | SVZ | 6.5 |
| 22 | Tumor mass | 0.896 |
| | SVZ | 0.469 |
| MIS1 | Tumor mass | 0.367 |
| | SVZ | 0.108 |
| MIS2 | Tumor mass | 7.37 |
| | SVZ | 6.03 |
| MIS3 | Tumor mass | 8.67 |
| | SVZ | 0.691 |
| Normal SVZ | HNS1 | 8.32 |
| | HNS2 | 2.68 |

**Supplementary Figure 1. Violin plots of the number of detected genes and UMI in each GBM patient and area.** Number of detected genes (top or left) and UMI (bottom or right) in each patient and each area: **a**, T_Mass; **b**, T_SVZ; and **c**, N_SVZ. UMI, unique molecular identifier.

**a**

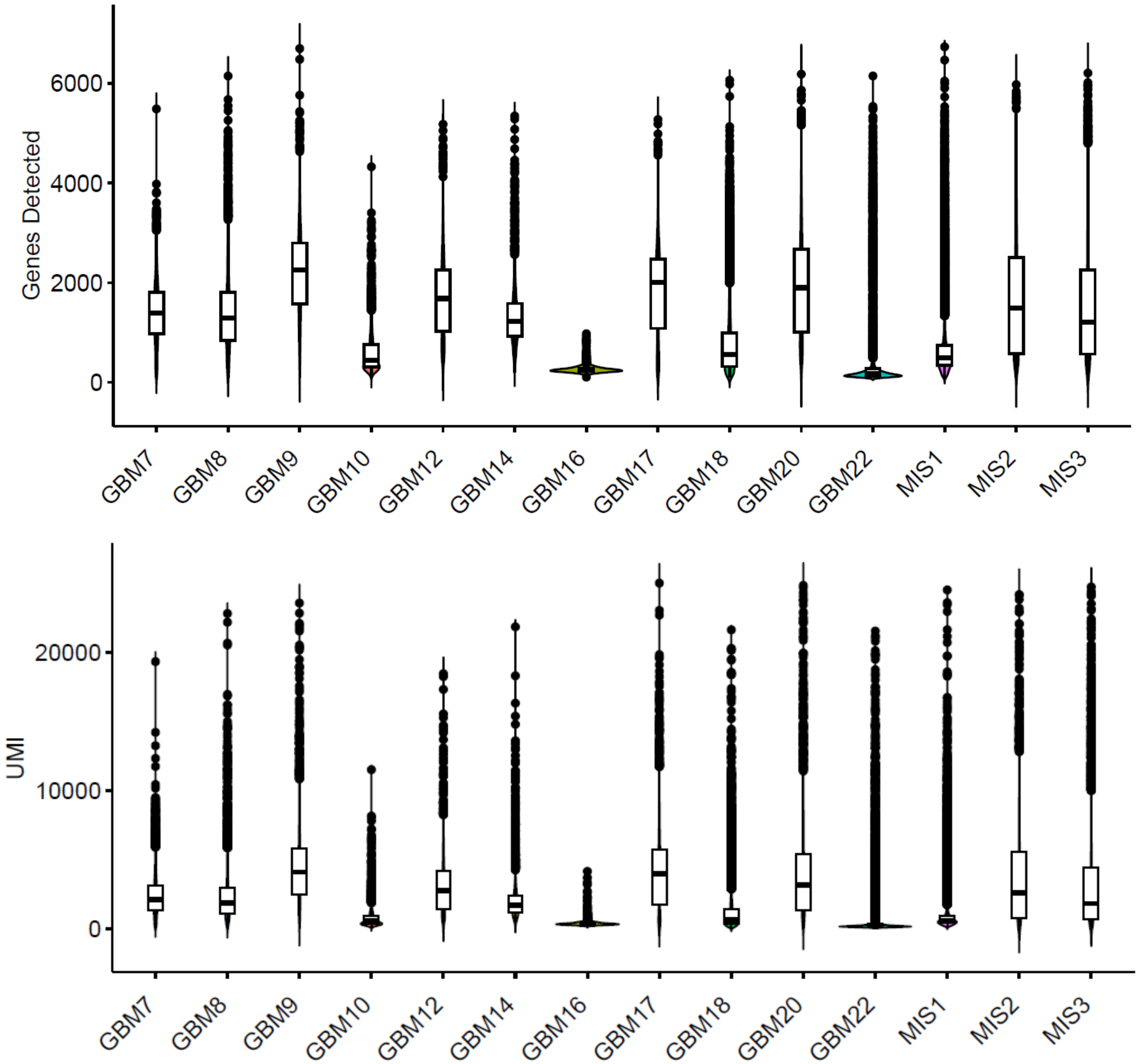

**b**

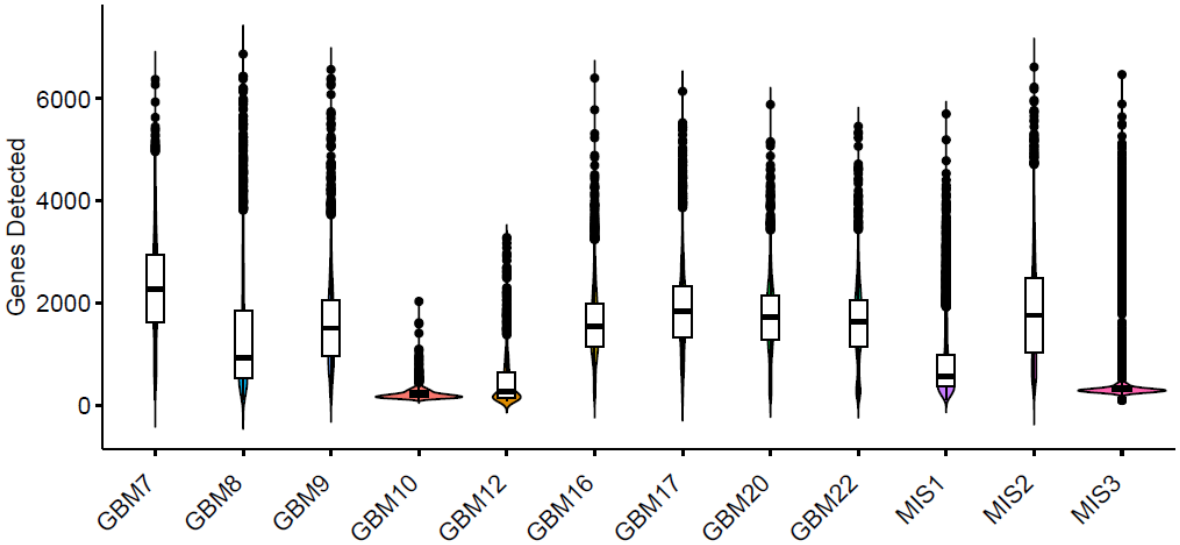

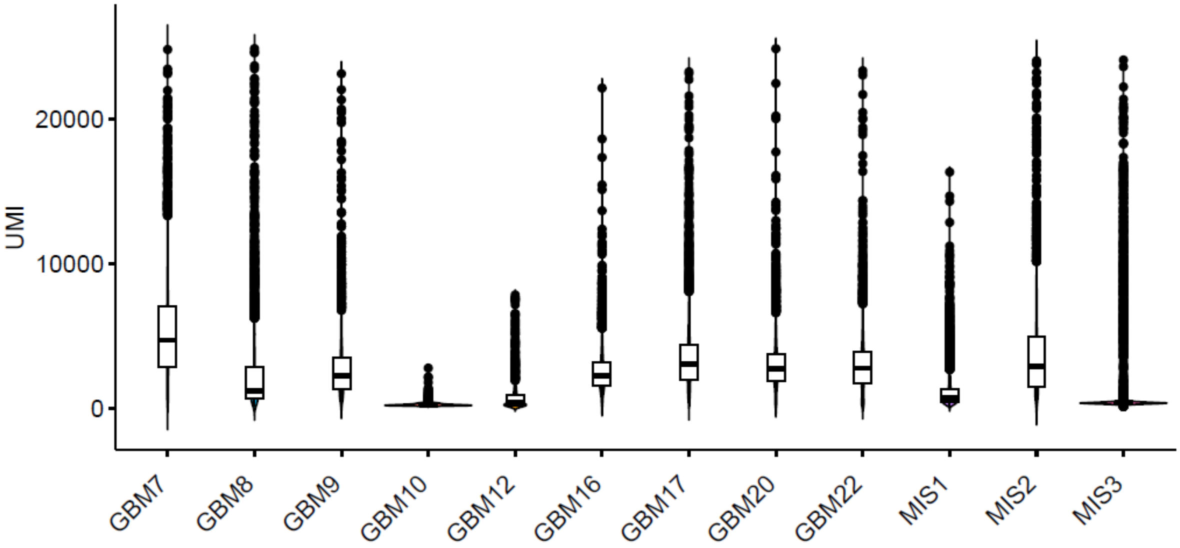

**c**

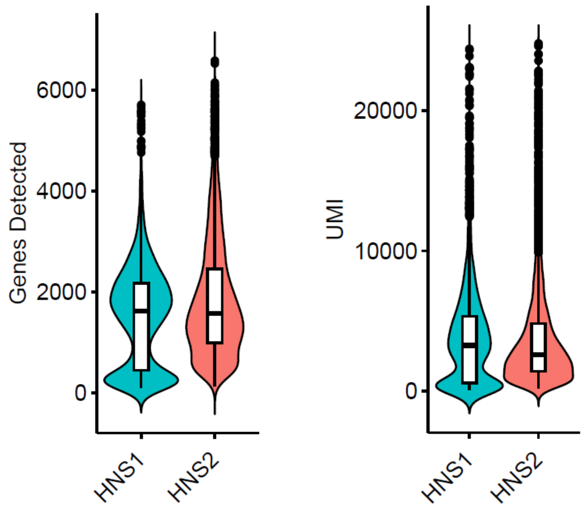

**Supplementary Figure 2. Copy-number variations for all GBM patients by area. a**, The reference cells from the integrated N_SVZ samples used in the copy-number variation analysis of the cancer cells of the integrated T_SVZ samples of Fig. 1e are shown here. **b, c, d,** Copy-number variations in the normal cells of the integrated T_SVZ samples (**b**), in the tumor cells of the integrated T_Mass samples (**c,** top and bottom) and in the normal cells of the integrated T_Mass samples (**d**) are shown on the y-axis. The genomic location of each variation is by chromosome (x-axis) and the reference cells used are from the integrated N_SVZ samples.

**a**

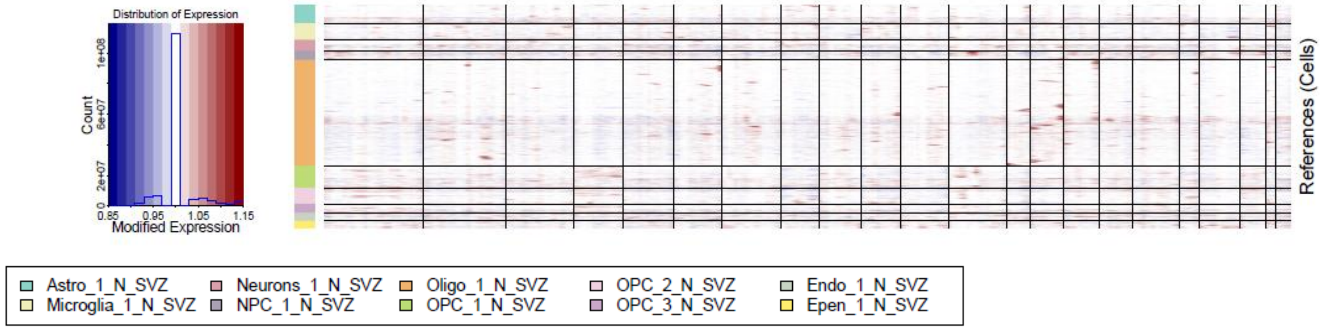

**b**

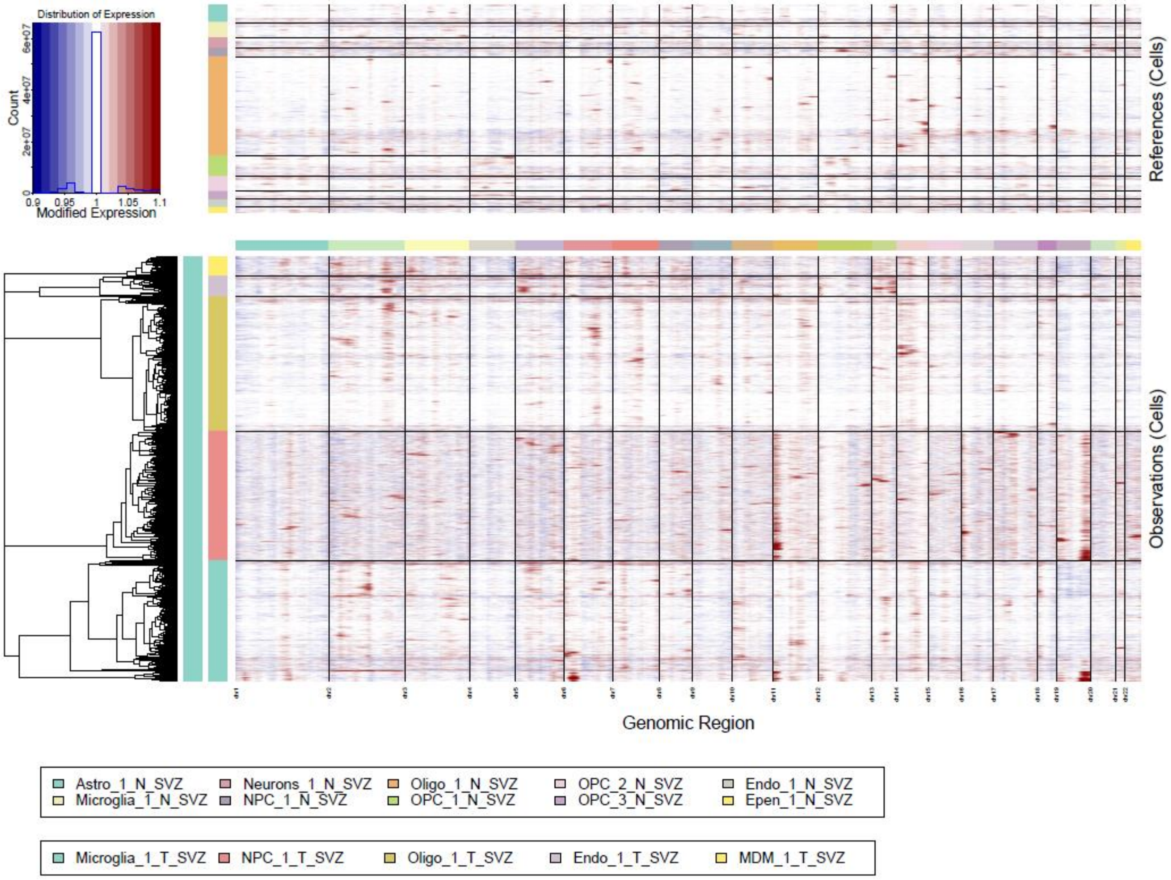
Distribution of Expression
Count
Modified Expression
References (Cells)
Observations (Cells)
Genomic Region
Astro_1_N_SVZ
Microglia_1_N_SVZ
Neurons_1_N_SVZ
NPC_1_N_SVZ
Oligo_1_N_SVZ
OPC_1_N_SVZ
OPC_2_N_SVZ
OPC_3_N_SVZ
Endo_1_N_SVZ
Epen_1_N_SVZ
Microglia_1_T_SVZ
NPC_1_T_SVZ
Oligo_1_T_SVZ
Endo_1_T_SVZ
MDM_1_T_SVZ

**c**

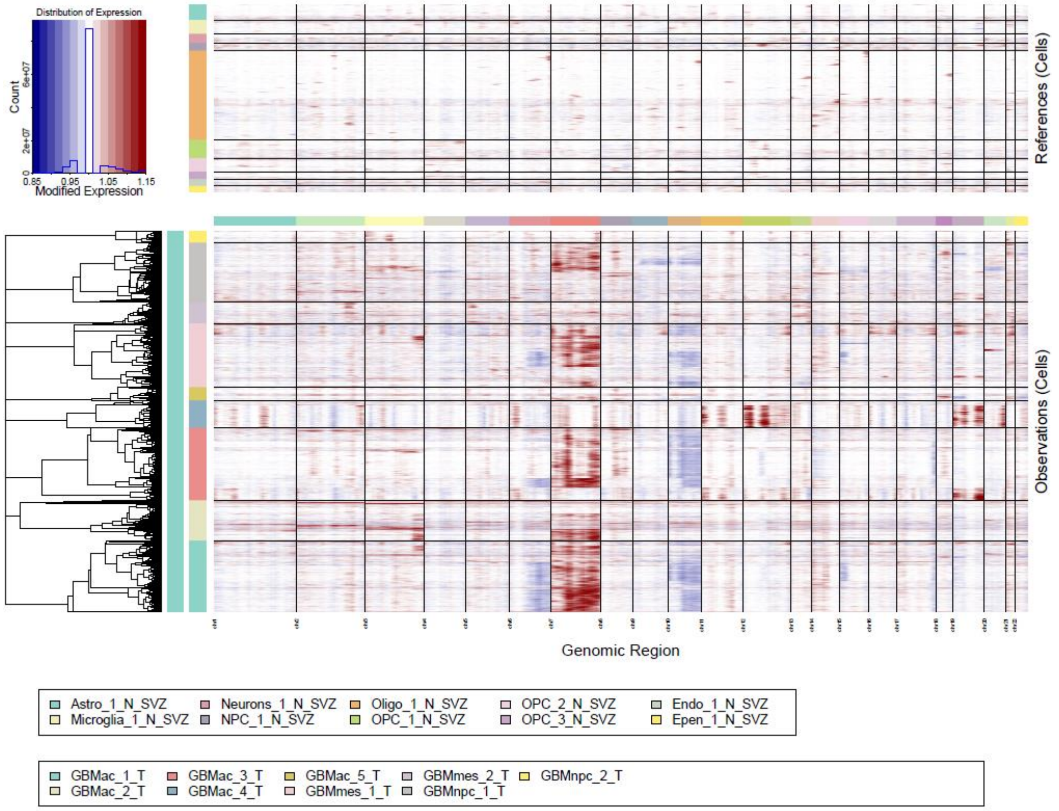
Distribution of Expression
Count
Modified Expression
References (Cells)
Observations (Cells)
Genomic Region
Astro_1_N_SVZ
Microglia_1_N_SVZ
Neurons_1_N_SVZ
NPC_1_N_SVZ
Oligo_1_N_SVZ
OPC_1_N_SVZ
OPC_2_N_SVZ
OPC_3_N_SVZ
Endo_1_N_SVZ
Epen_1_N_SVZ
GBMac_1_T
GBMac_2_T
GBMac_3_T
GBMac_4_T
GBMac_5_T
GBMmes_1_T
GBMmes_2_T
GBMnpc_1_T
GBMnpc_2_T

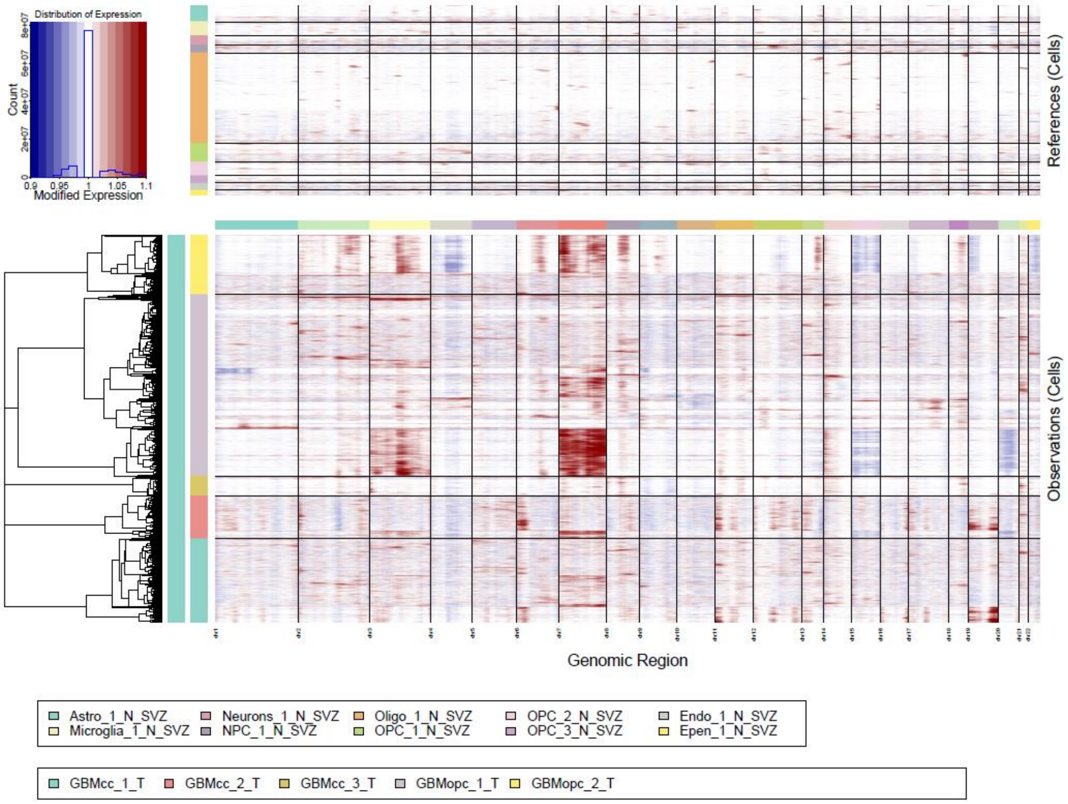
Distribution of Expression
Count
Modified Expression
References (Cells)
Observations (Cells)
Genomic Region
Astro_1_N_SVZ
Microglia_1_N_SVZ
Neurons_1_N_SVZ
NPC_1_N_SVZ
Oligo_1_N_SVZ
OPC_1_N_SVZ
OPC_2_N_SVZ
OPC_3_N_SVZ
Endo_1_N_SVZ
Epen_1_N_SVZ
GBMcc_1_T
GBMcc_2_T
GBMcc_3_T
GBMopc_1_T
GBMopc_2_T

**d**

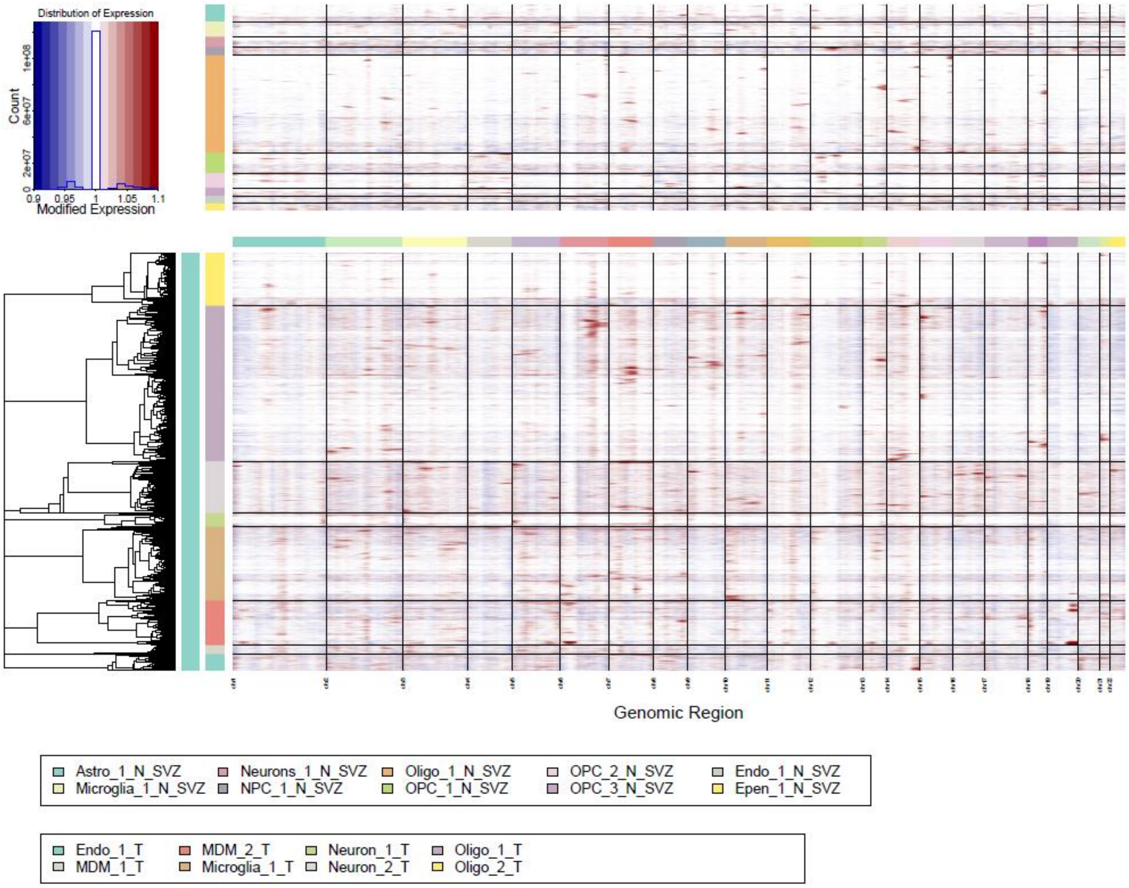
Distribution of Expression
Count
Modified Expression
References (Cells)
Observations (Cells)
Genomic Region
Astro_1_N_SVZ
Microglia_1_N_SVZ
Neurons_1_N_SVZ
NPC_1_N_SVZ
Oligo_1_N_SVZ
OPC_1_N_SVZ
OPC_2_N_SVZ
OPC_3_N_SVZ
Endo_1_N_SVZ
Epen_1_N_SVZ
Endo_1_T
MDM_1_T
MDM_2_T
Microglia_1_T
Neuron_1_T
Neuron_2_T
Oligo_1_T
Oligo_2_T

**Supplementary Figure 3. Cell type annotations using three classifiers.** Heatmaps of the enrichment scores for each cluster of the integrated T_Mass samples (**a**), T_SVZ samples (**b**), and N_SVZ samples (**c**). For the T_Mass and the T_SVZ, the following classifiers are shown from left to right: the pan-glioma by Johnson *et al.*[2] and the pathway-based by Garofano *et al.*[3], both as differentially expressed genes (DE) and all genes, and the cell state classification by Neftel *et al.*[4]. For the N_SVZ samples, the marker genes used for cell annotation were taken from McKenzie *et al.*[5].

**a**

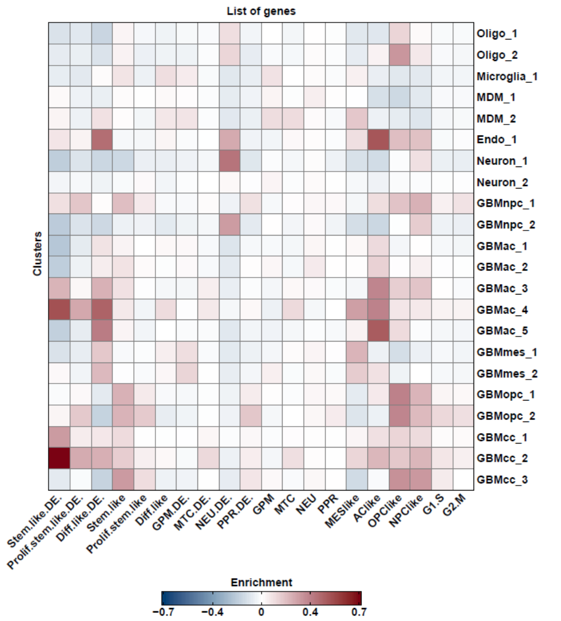

**b**

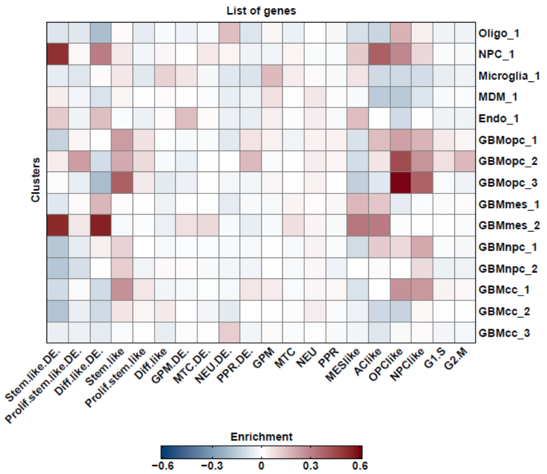

**c**

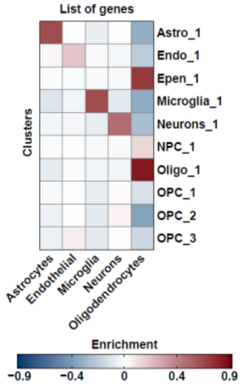

**Supplementary Figure 4. Tumor cell state proportions in each patient area.** Bar graphs showing the percentage of each cancer cell state[4] in each patient area: **a**, T_Mass; **b**, T_SVZ.

**a**

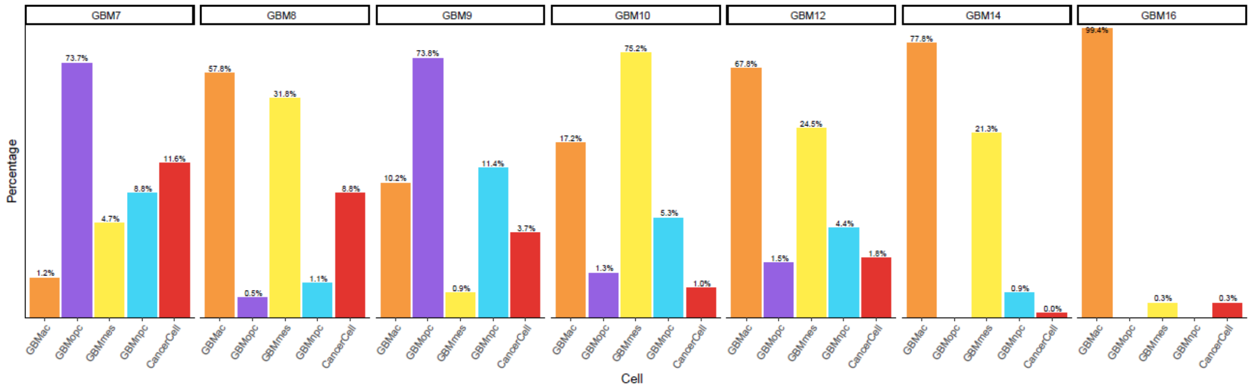

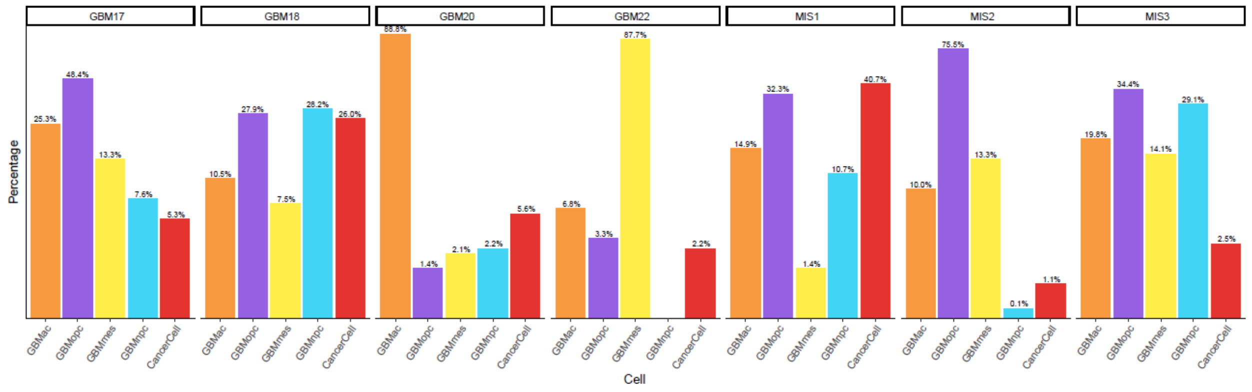

**b**

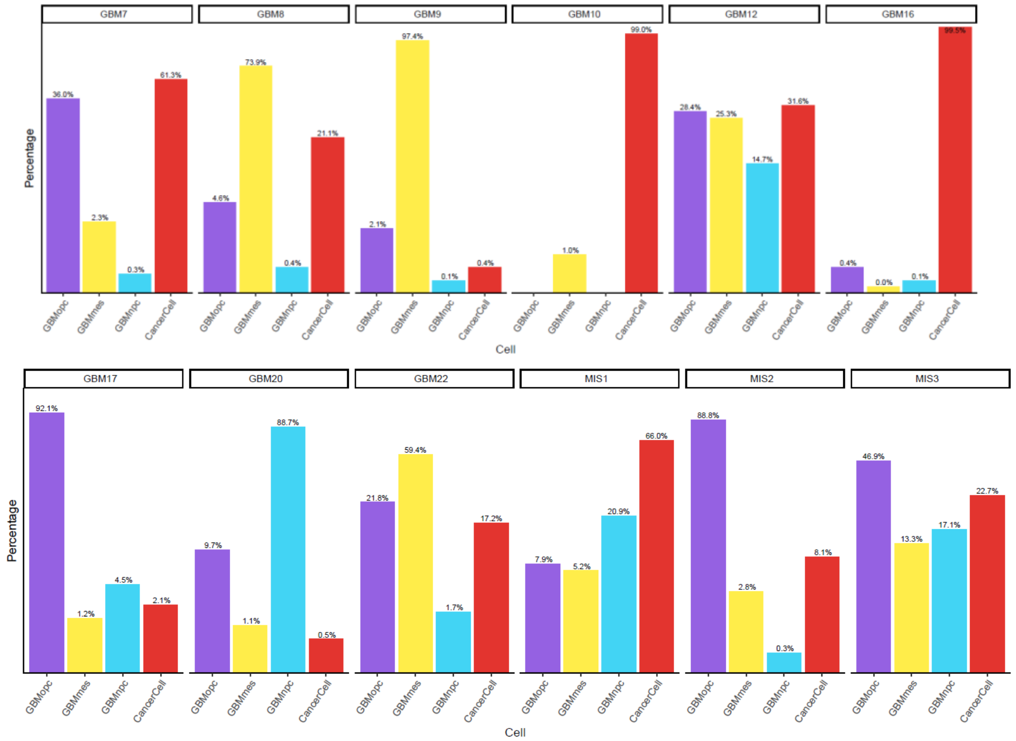

**Supplementary Figure 5. Initial and terminal macrostates of the tumor cells in the T_Mass and the T_SVZ.** CellRank-computed initial and terminal macrostates in the T_Mass (**a**) and the T_SVZ (**b**). The macrostates in each area are color-coded as in Fig. 2a.

**a**

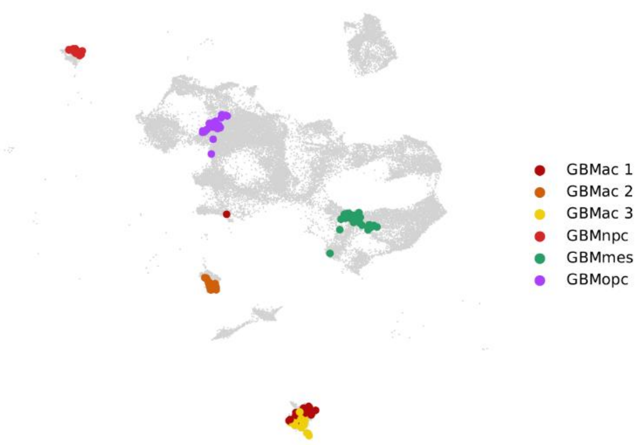

**b**

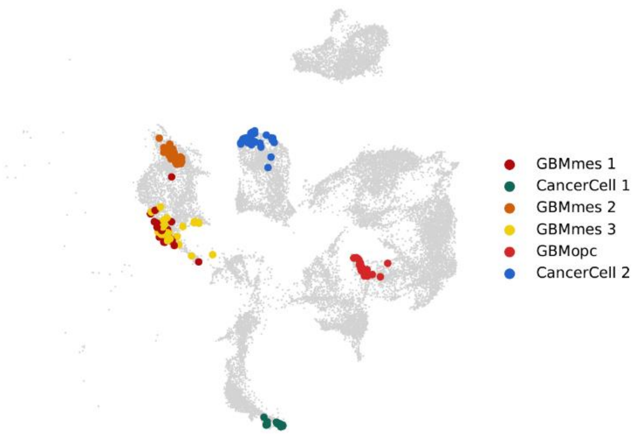

**Supplementary Figure 6. Fate probabilities of the tumor cells in the T_Mass and the T_SVZ.** CellRank-computed fate probabilities in the T_Mass (**a**) and the T_SVZ (**b**). The same color code of Fig. 2a is used here.

**a** **b**

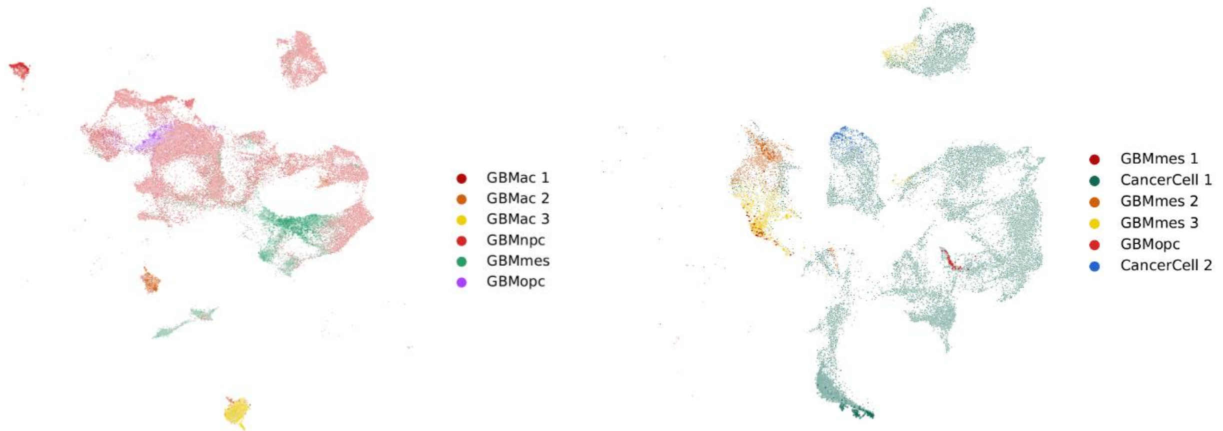

**Supplementary Figure 7. Differentially expressed genes between the T_Mass and the N_SVZ.** Volcano plots showing the differentially expressed genes in the comparisons between the T_Mass and the N_SVZ as whole areas, left, and as microglia only, right. In all the analyses, average log2(Fold Change)>0.3 and $p$<0.05 were used.

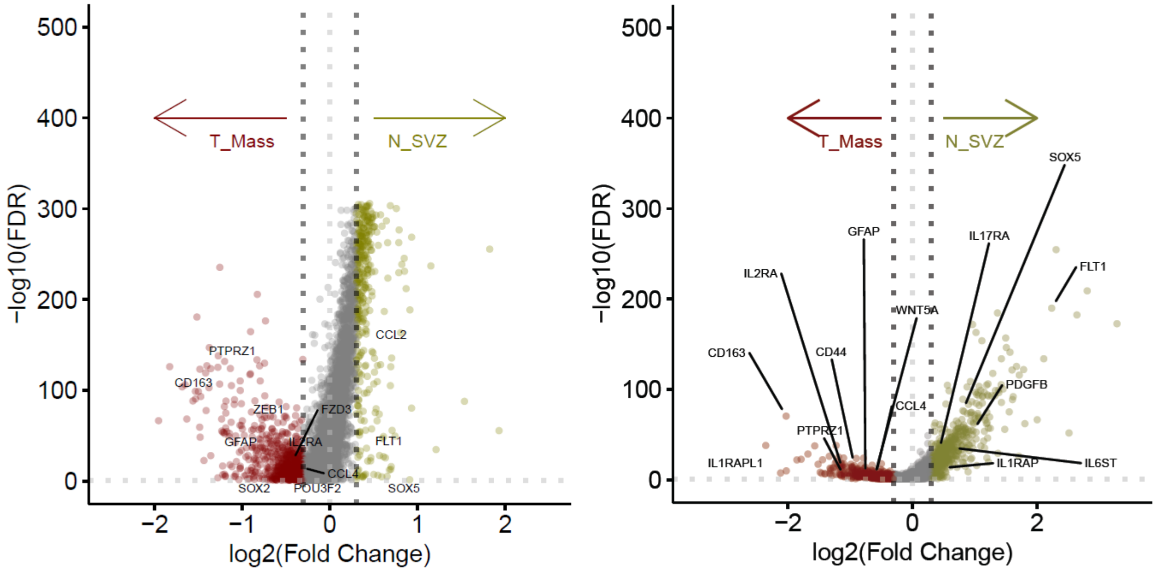

**Supplementary Table 4. List of differentially expressed genes between N_SVZ *versus* T_Mass, N_SVZ *versus* T_SVZ and T_Mass *versus* T_SVZ.** This table includes a list of all the genes differentially expressed in the comparisons between T_Mass, T_SVZ and N_SVZ. The average log2Fold Change (avg_logFC), $p$ value, adjusted $p$ value and direction of differential expression in each comparison are also included. This table is included as Appendix 1.

**Supplementary Table 5. List of differentially expressed genes in microglia between N_SVZ *versus* T_Mass, N_SVZ *versus* T_SVZ and T_Mass *versus* T_SVZ.** This table includes a list of all the genes differentially expressed in microglia in the comparisons between T_Mass, T_SVZ and N_SVZ. The average log2Fold Change (avg_logFC), $p$ value, adjusted $p$ value and direction of differential expression in each comparison are also included. This table is included as Appendix 2.

**Supplementary Figure 8. Gene expression signature of 'homeostatic', 'activated', and 'inflammatory' microglia in the N_SVZ, T_Mass, and T_SVZ.** Heatmap of gene expression signature associated with 'homeostatic', 'activated', and 'inflammatory' microglia in the three areas.

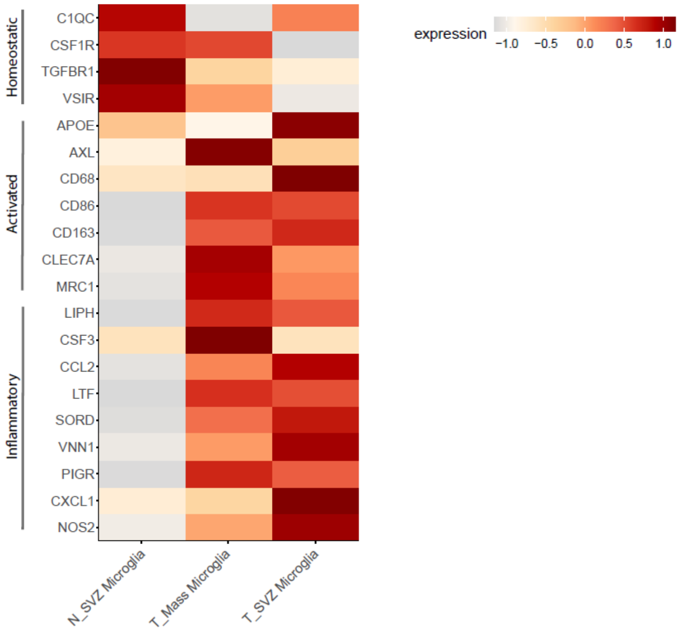

**Supplementary Figure 9. Tumor cell purity in the T_Mass and the T_SVZ of the 4 GBM patients and in the HNS1 sample.** Bar graph showing the proportion of tumor (blue) and normal (orange) cells in each area (T_Mass and T_SVZ) of the 4 GBM patients and in the HNS1 sample.

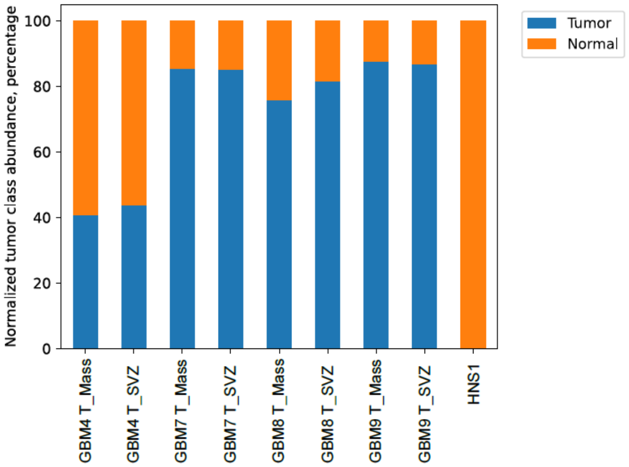

**Supplementary Figure 10. Cell type composition in the T_Mass and the T_SVZ of the 4 GBM patients and in the HNS1 sample.** Bar graphs showing the proportion of each cell type in each area (T_Mass and T_SVZ) of the 4 GBM patients and in the HNS1 sample. Tumor cells are annotated using the cell state classification by Neftel *et al.*[4].

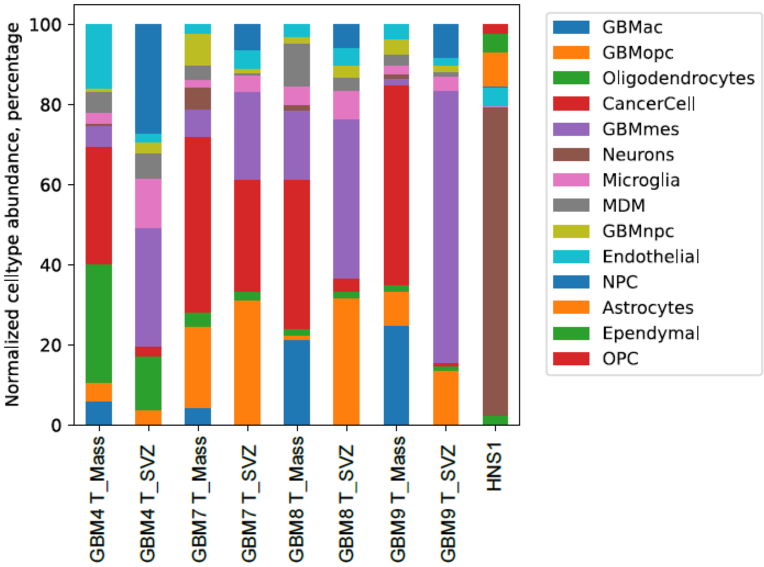

**Supplementary Figure 11. Spatial frequency correlation between microglia and all the other cell types in the T_Mass and the T_SVZ.** Cell frequency correlation graphs of microglia and tumor and normal cells in the T_Mass and the T_SVZ of each patient: **a**, GBM4 T_Mass; **b**, GBM4 T_SVZ; **c**, GBM7 T_Mass; **d**, GBM7 T_SVZ; **e**, GBM8 T_Mass; **f**, GBM8 T_SVZ; **g**, GBM9 T_Mass; **h**, GBM9 T_SVZ; **i**, HNS1. Linear regression is shown on each graph.

**a**

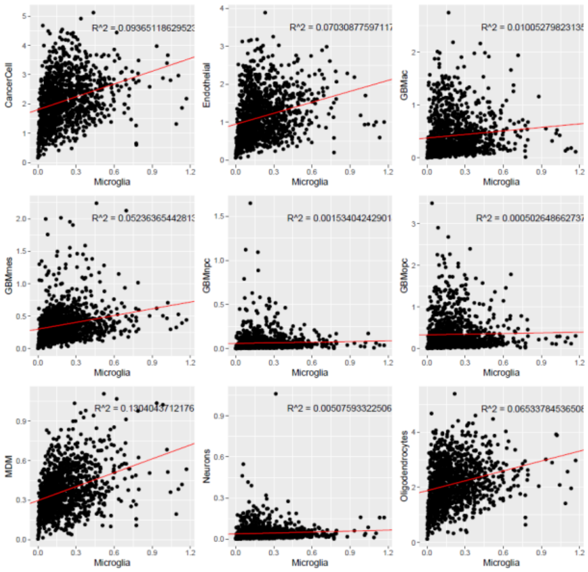

**b**

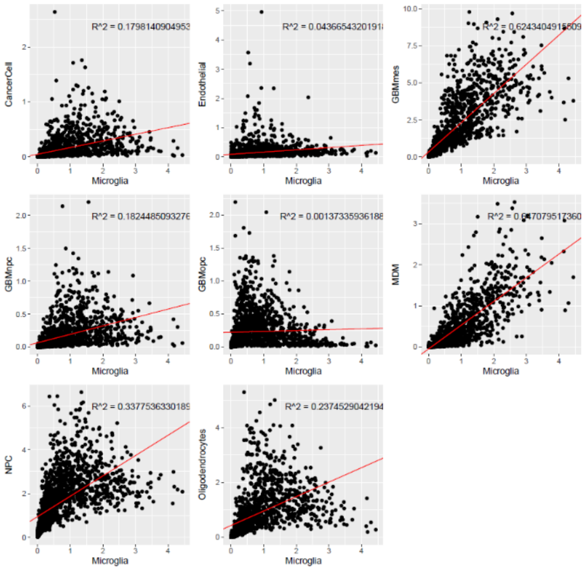

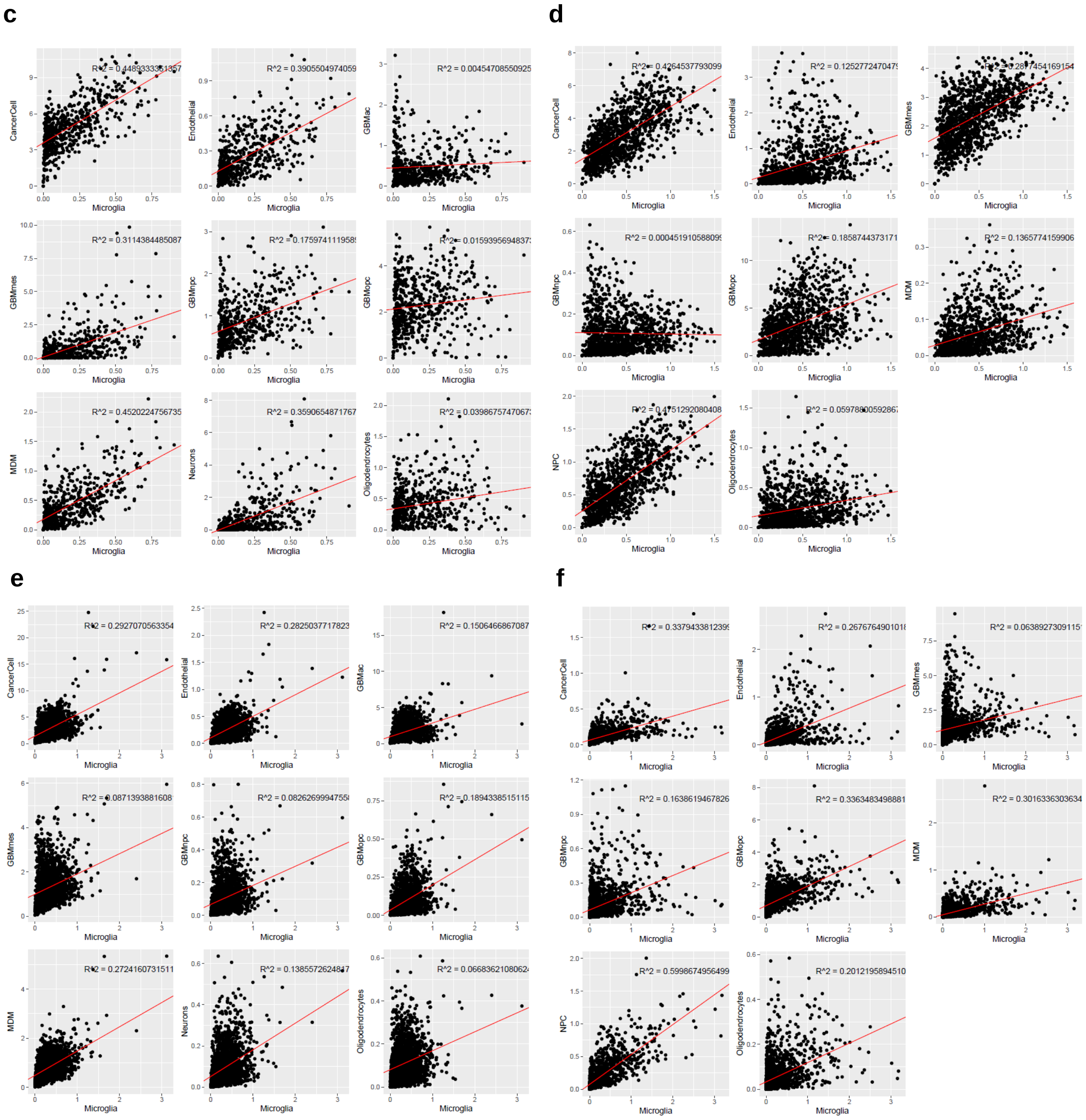
c
CancerCell
R^2 = 0.448333335185
Endothelial
R^2 = 0.3905504974059
GBMac
R^2 = 0.00454708550925
GBMmes
R^2 = 0.3114384485087
GBMnpc
R^2 = 0.175974111958
GBMopc
R^2 = 0.0159395694837
MDM
R^2 = 0.4520224756735
Neurons
R^2 = 0.3590654871767
Oligodendrocytes
R^2 = 0.0398675747067
Microglia
d
CancerCell
R^2 = 0.4264537793099
Endothelial
R^2 = 0.125277247047
GBMmes
R^2 = 0.2877454169154
GBMnpc
R^2 = 0.000451910588099
GBMopc
R^2 = 0.1858744373171
MDM
R^2 = 0.1365774159906
NPC
R^2 = 0.4751292080408
Oligodendrocytes
R^2 = 0.0597860059286
Microglia
e
CancerCell
R^2 = 0.2927070563354
Endothelial
R^2 = 0.2825037717823
GBMac
R^2 = 0.1506466867087
GBMmes
R^2 = 0.0871393881608
GBMnpc
R^2 = 0.0826269994755
GBMopc
R^2 = 0.1894338515115
MDM
R^2 = 0.2724160731511
Neurons
R^2 = 0.138557262484
Oligodendrocytes
R^2 = 0.0668362108062
Microglia
f
CancerCell
R^2 = 0.337943381239
Endothelial
R^2 = 0.267676490101
GBMmes
R^2 = 0.063892730911
GBMnpc
R^2 = 0.1638619467826
GBMopc
R^2 = 0.3363483498881
MDM
R^2 = 0.3016336303634
NPC
R^2 = 0.5998674956499
Oligodendrocytes
R^2 = 0.2012195894510
Microglia

g

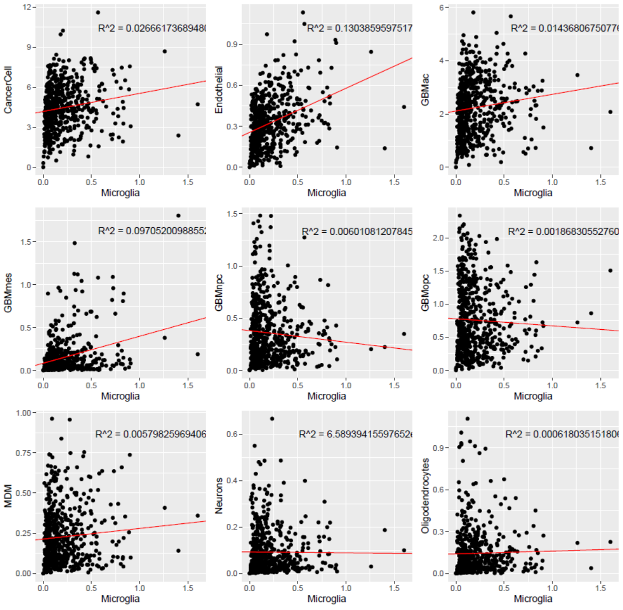

h

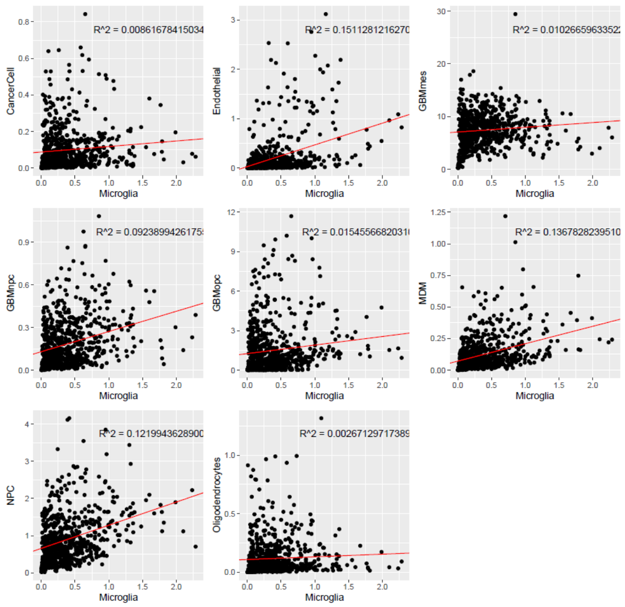

i

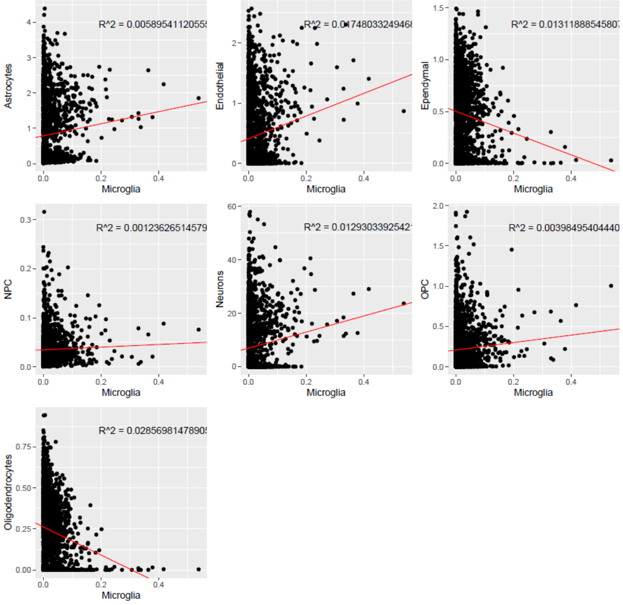

**Supplementary Figure 12. Patterns of spatial dependencies among cell types in the T_Mass and T_SVZ of the 4 GBM patients and in the HNS1 sample.** Type coupling analysis of spatial dependencies among cell types in the T_Mass and the T_SVZ of each patient: GBM4 T_Mass (**a**) and T_SVZ (**b**); GBM7 T_Mass (**c**) and T_SVZ (**d**); GBM8 T_Mass (**e**) and T_SVZ (**f**); GBM9 T_Mass (**g**) and T_SVZ (**h**); HNS1 (**i**). The edge width is proportional to the L1 norm of the vector of fold changes of differentially expressed genes for each pair of sender and receiver cell types and only edges with at least 150 genes are visualized.

a

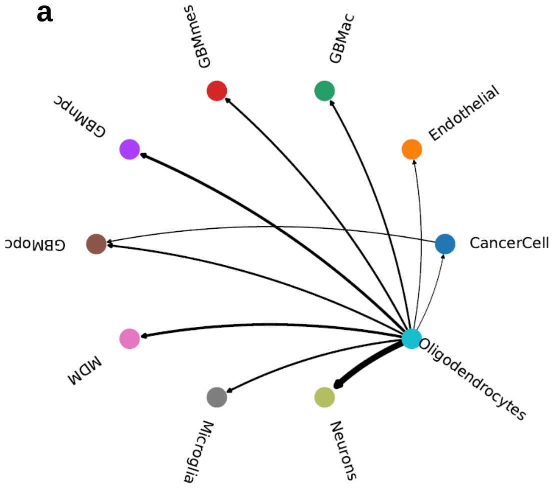

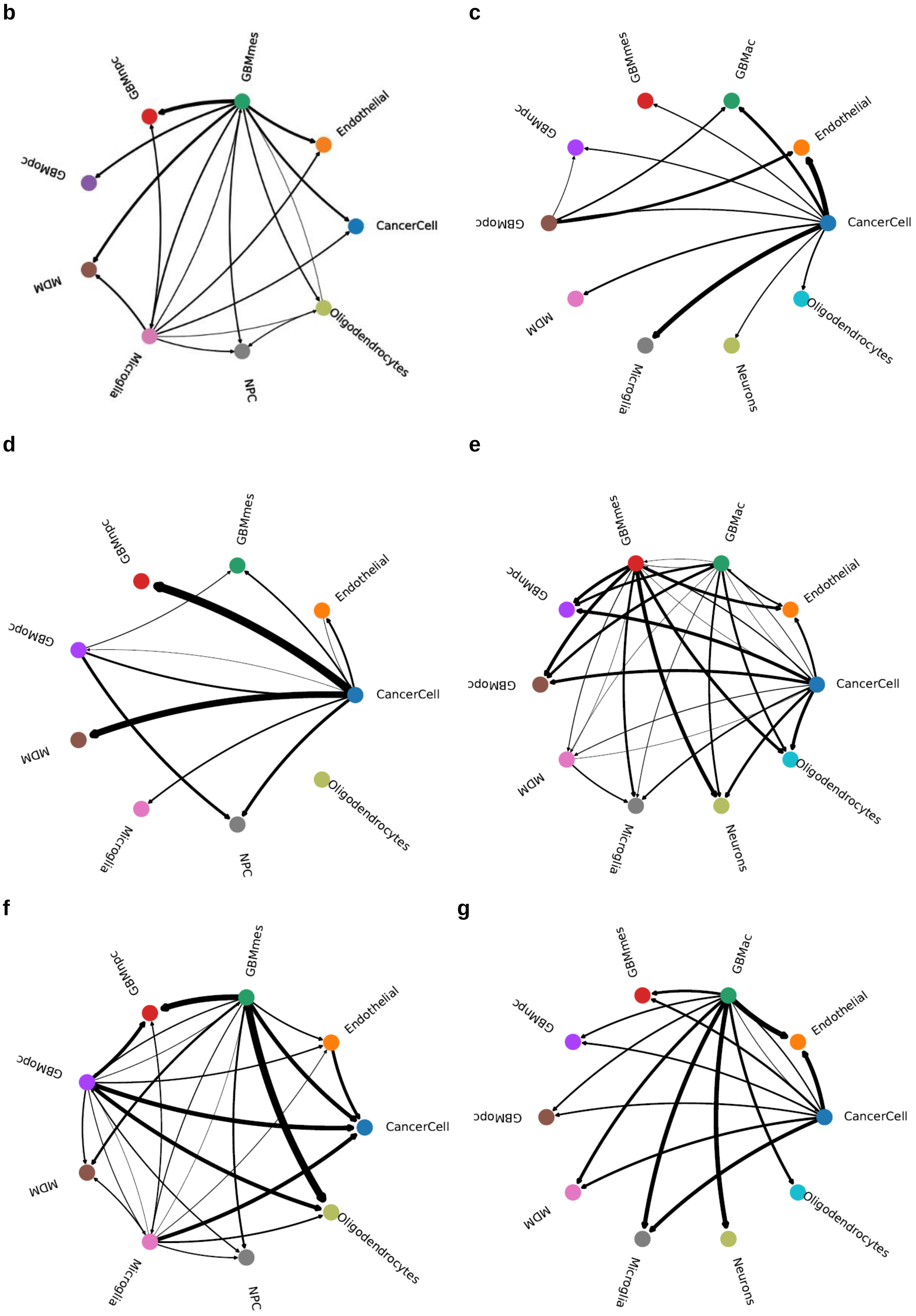
b
GBMmes
GBMnpc
Endothelial
GBMopc
CancerCell
MDM
Oligodendrocytes
Microglia
NPC
c
GBMmes
GBMac
GBMnpc
Endothelial
GBMopc
CancerCell
MDM
Oligodendrocytes
Microglia
Neurons
d
GBMnpc
GBMmes
Endothelial
GBMopc
CancerCell
MDM
Oligodendrocytes
Microglia
NPC
e
GBMmes
GBMac
GBMnpc
Endothelial
GBMopc
CancerCell
MDM
Oligodendrocytes
Microglia
Neurons
f
GBMnpc
GBMmes
Endothelial
GBMopc
CancerCell
MDM
Oligodendrocytes
Microglia
NPC
g
GBMmes
GBMac
GBMnpc
Endothelial
GBMopc
CancerCell
MDM
Oligodendrocytes
Microglia
Neurons

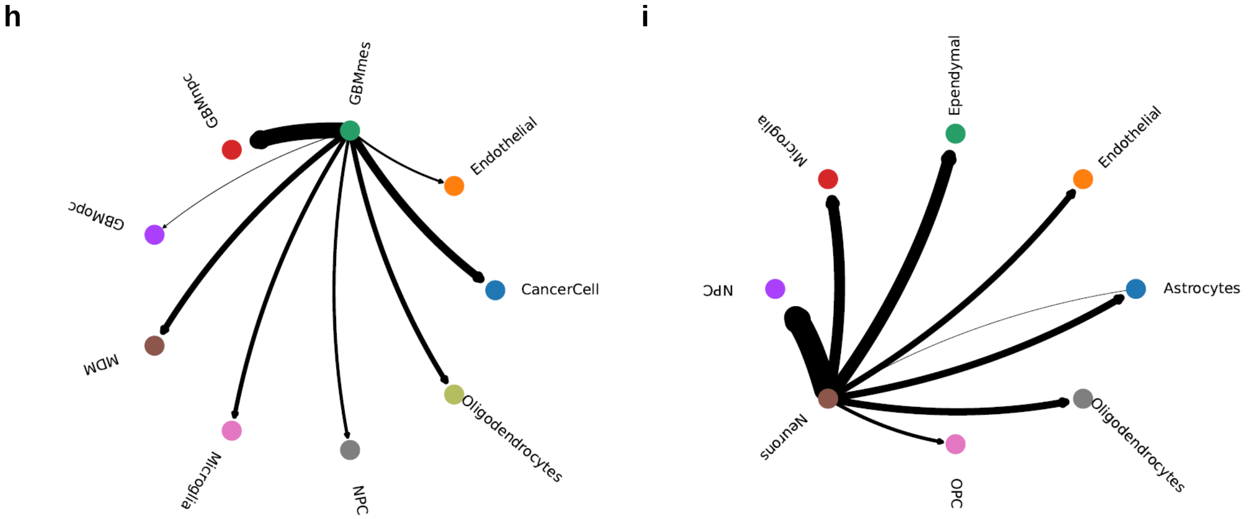

**Supplementary Figure 13. Inferred interactions in the N_SVZ, T_SVZ, and T_Mass.** Bar graph showing the number of inferred interactions in the N_SVZ, T_SVZ, and T_Mass.

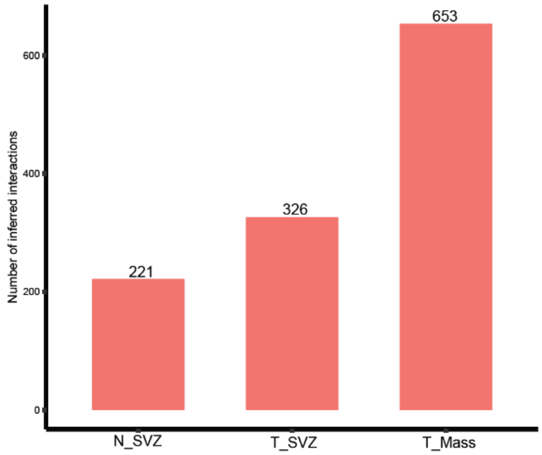

**Supplementary Figure 14. Predicted incoming signaling patterns in the T_Mass, the T_SVZ and the N_SVZ.** Heatmaps of incoming signaling pathways in the T_Mass (left), T_SVZ (middle) and N_SVZ (right).

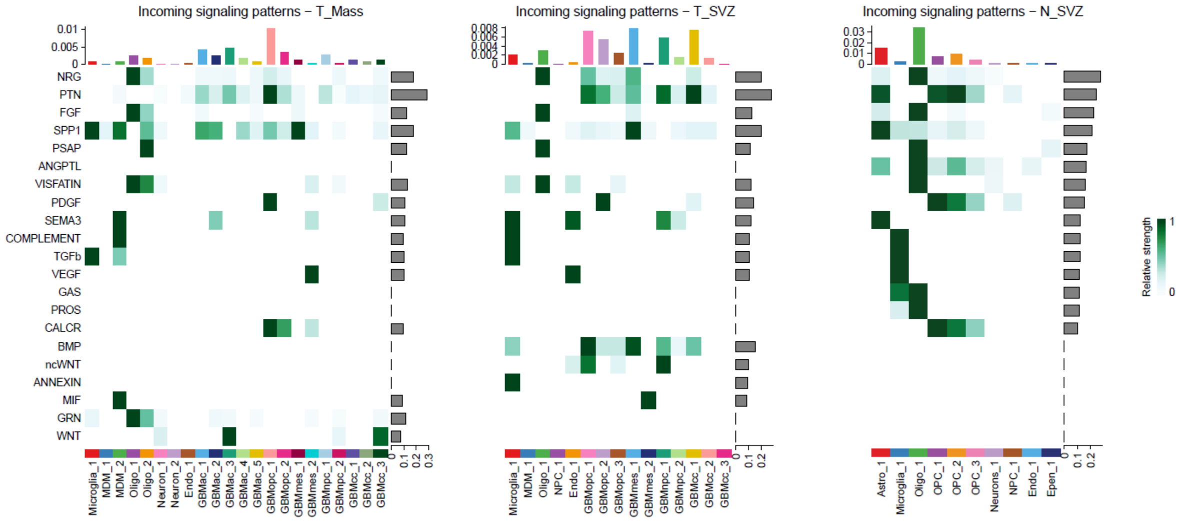

**Supplementary Figure 15. Ligand-receptor prediction analysis between microglia/MDM and any other cell cluster of the T_Mass and the N_SVZ.** Dot plots of the ligand-receptor prediction analysis between microglia (in **a**) and the two MDM clusters (in **b**, MDM_1, left and MDM_2, right) and any other cell cluster of the T_Mass. **c**, the same analysis was also performed between microglia and any other cell cluster of the N_SVZ.

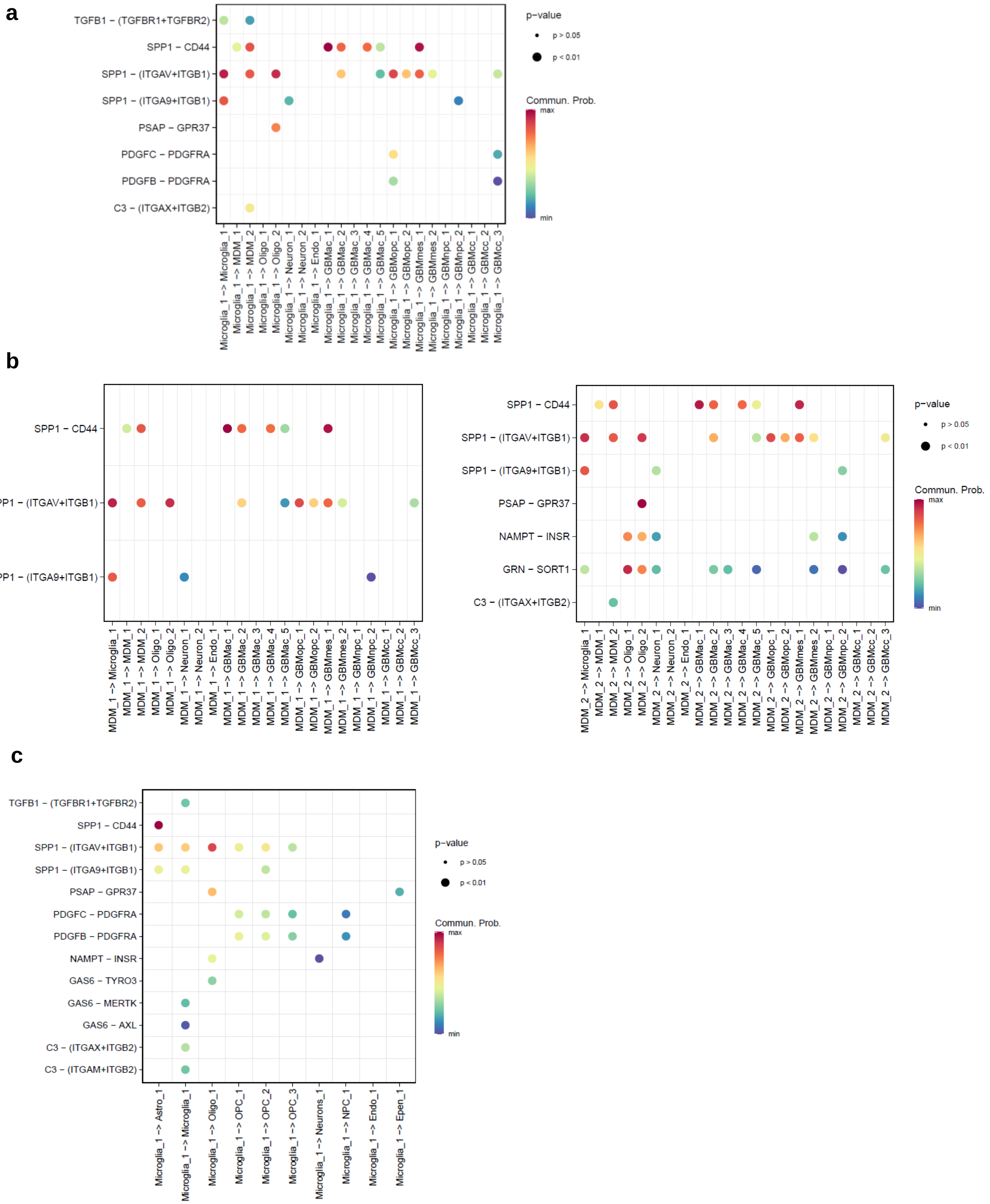

**Supplementary Figure 16. Cell type expression of *WNT5A*, *IL1B*, *FZD3*, and *IL1RAP* in the T_Mass and N_SVZ.** Dot plots showing the cell type expression of *WNT5A*, *IL1B*, *FZD3*, and *IL1RAP* in the T_Mass (left) and N_SVZ (right).

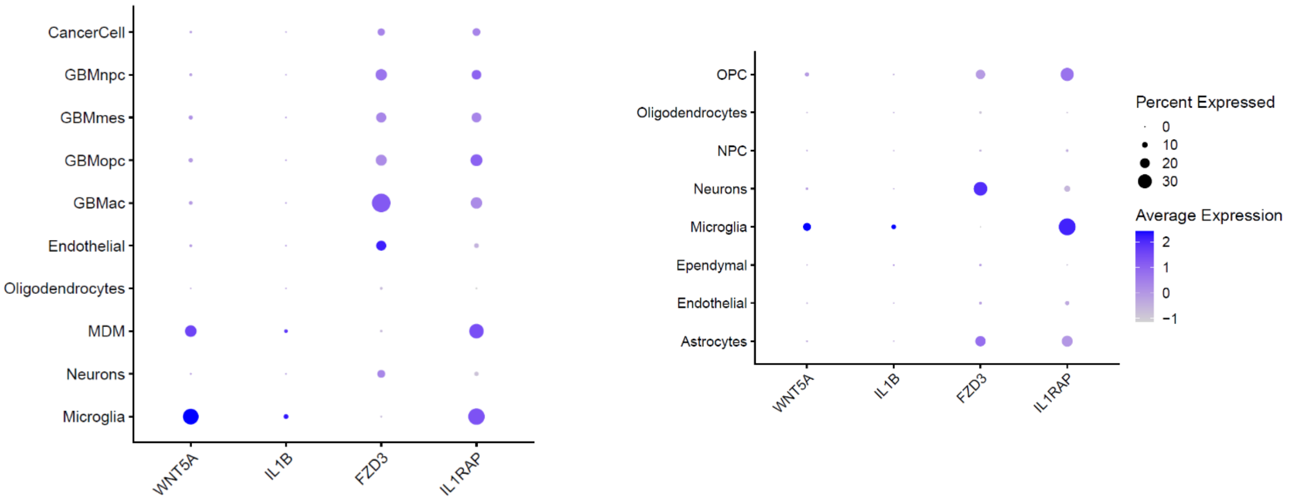

**Supplementary Figure 17. IL1-RAcP immunofluorescence of TAMs isolated from the T_Mass and the T_SVZ.** Representative images of TAMs stained for IL1-RAcP and counterstained with DAPI. The TAMs were isolated from GBM7. Scale bars, 50 µm.

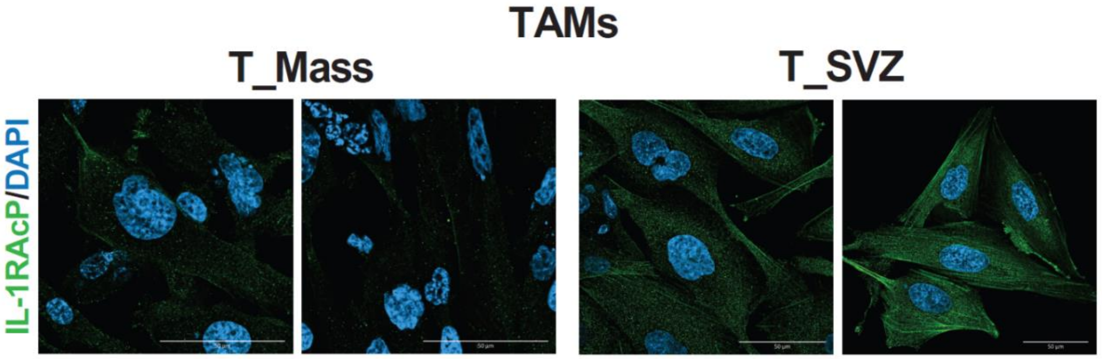

**Supplementary Figure 18. Transwell assays of CSCs isolated from the T_Mass and the T_SVZ of GBM17 and exposed to conditioned medium of matched TAMs treated with Box5.** Quantification of cell migration based on absorbance of eluted crystal violet used in transwell assays of CSCs. ****=p<0.0001.

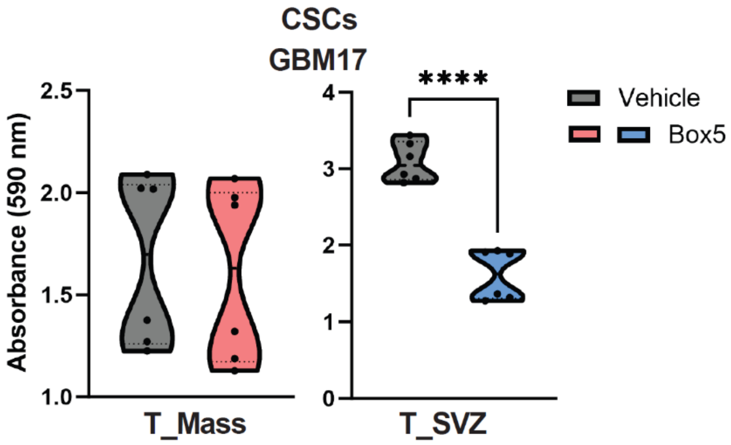

### References

1 Verhaak, R. G. *et al.* Integrated genomic analysis identifies clinically relevant subtypes of glioblastoma characterized by abnormalities in PDGFRA, IDH1, EGFR, and NF1. *Cancer Cell* **17**, 98-110, doi:10.1016/j.ccr.2009.12.020 (2010).

2 Johnson, K. C. *et al.* Single-cell multimodal glioma analyses identify epigenetic regulators of cellular plasticity and environmental stress response. *Nat Genet* **53**, 1456-1468, doi:10.1038/s41588-021-00926-8 (2021).

3 Garofano, L. *et al.* Pathway-based classification of glioblastoma uncovers a mitochondrial subtype with therapeutic vulnerabilities. *Nat Cancer* **2**, 141-156, doi:10.1038/s43018-020-00159-4 (2021).

4 Neftel, C. *et al.* An Integrative Model of Cellular States, Plasticity, and Genetics for Glioblastoma. *Cell* **178**, 835-849 e821, doi:10.1016/j.cell.2019.06.024 (2019).

5 McKenzie, A. T. *et al.* Brain Cell Type Specific Gene Expression and Co-expression Network Architectures. *Sci Rep* **8**, 8868, doi:10.1038/s41598-018-27293-5 (2018).
